# Integrated metabolomics and proteomics from voxelated cortical hemispheres of adult rhesus monkeys

**DOI:** 10.64898/2026.04.04.716423

**Authors:** Qiuyi Wu, Alev M. Brigande, Michael W. Lutz, Pixu Shi, Anita A. Disney

**Author notes:** contributed equally (Qiuyi Wu, Alev M. Brigande).

## Abstract

The spatial organization of molecular networks across cortex likely contributes to differences in local circuit vulnerability in aging and Alzheimer’s disease; yet many existing molecular datasets sacrifice spatial structure, sampling only a handful of regions per brain. Here, we present a framework for generating spatially registered, paired metabolomic and proteomic maps across an entire cortical hemisphere of an adult rhesus monkey, at millimeter resolution. One hemisphere each from two animals was harvested under controlled conditions, approximately flattened, and hand dissected at different sampling resolutions (roughly 2.5 and 4 mm/side) into tissue voxels. Each voxel was split after homogenization and extraction to provide matched aliquots for targeted metabolomics and deep untargeted proteomics. To handle these high dimensional data, we developed PChclust, a principal component guided feature clustering algorithm. For cross omic integration, we developed a spatially regularized sparse canonical correlation analysis (sr-sCCA), which incorporates spatial neighborhood structure via graph Laplacian smoothing. We recover meaningful biology: Molecular similarity between neighboring voxels decayed with distance in both modalities, confirming that voxelation captures spatially organized biological variance. The sr-sCCA identified joint proteome-metabolome components with coherent cortical gradients that were conserved across animals. Pathway enrichment analysis recovered brain relevant ontologies and reconstructed complete metabolic circuits from single voxels.

## Introduction

Tools and model systems for studying the prodromal phase of Alzheimer’s disease (AD), particularly its most common spontaneous, or late onset, form (LOAD), are lacking. The central difficulty in the study of LOAD^1^ is not a lack of molecular hypotheses; it is that many hypotheses are underdetermined by the measurements we typically make. This underdetermination, in part, reflects three difficult facts: pathology begins ∼20 years before diagnosis^2–7^; pathology and function are not tightly coupled^8^; and the paths to pathology are many. These observations imply that comparisons between diagnosed individuals and matched controls can be both reproducible, and mechanistically ambiguous: they pool individuals who may share a diagnostic label but not a causal route, and they often sample tissue after much of the relevant biology has already unfolded.

One response to these challenges is to change the unit of inference. A spatially resolved molecular case study within one individual has a key advantage: at a fixed time point, risk factors and insults are held constant, and regional differences can be interpreted against that shared background. Spatially resolved studies at the scale of the whole cortex, in particular, are likely to be important because many LOAD-associated pathologies follow predictable spatiotemporal patterns^9–11^. If these spatial patterns reveal regional differences in vulnerability or resilience, then measurements that erase spatial structure discard relevant information.

Despite this potential, most molecular datasets in Alzheimer’s disease neuroscience are spatially coarse. We typically sample a handful of regions, homogenize each, and search for associations with clinical or pathological labels. This strategy cannot represent gradients, boundaries, or patchy structure across the cortical sheet, and makes it difficult to distinguish true biological differences from differences in cellular composition, vascular content, or local microenvironment.

Here, we present a proof-of-concept framework that enables spatially resolved molecular comparisons across the cortex, using paired metabolomics and proteomics. Our underlying motivation is to work toward understanding the LOAD prodrome, but the framework is general: it produces spatially registered molecular maps that can be aligned to pathology, cytoarchitecture, and connectivity. Deploying this method towards understanding LOAD and other pathologies will require hypothesis-driven analyses of within-subjects datasets such as these, comparative case studies, and population comparisons. Enabling those studies is our goal.

Why the cortex of rhesus monkeys? Many LOAD-relevant vulnerability patterns are properties of cortex^9,12^, which is a structured sheet embedded with primate-specific association networks^13^. Sufficiently detailed spatial mapping is difficult to obtain from humans — particularly in the prodromal phase — because the relevant individuals are hard to identify pre-mortem, documentation of environmental factors is limited, and preservation of biomolecules is challenging under typical harvest conditions (see Results, and^14,15^). In rhesus monkeys, tissue collection can be controlled in ways that reduce these confounds, and animals can be studied at ages corresponding to the human prodromal window, as we do here.

Why paired metabolomics and proteomics? The biological questions motivating LOAD research — energetics, oxidative stress, lipid remodeling, neurotransmitter metabolism, proteostasis — are pathway questions. The proteome reflects catalytic and structural capacity; the metabolome reflects biochemical activity at the time of measurement; either layer alone is easy to over-interpret^16–18^. Paired measurements constrain interpretation by asking whether a pathway-level narrative has support on both sides of the enzyme–substrate divide.

Spatial multi-omics always involves a three-way trade between spatial resolution, molecular coverage, and molecular depth. Here, we provide a harvest and mass spectrometry analysis procedure that makes it feasible to generate spatially registered, paired multi-omic cortical maps at a scale appropriate for asking vulnerability questions in large brains. We describe the voxelation and reconstruction procedure, and the metabolomics and proteomics pipelines and quality control strategy. Dense sampling creates a statistical challenge: neighboring samples are not independent, and standard inference ignores this structure. We therefore also provide a novel analysis framework for dimensionality reduction and spatially informed integration. Finally, we demonstrate that the resulting data support interpretable biological structure at the pathway level. The larger claim is pragmatic: if you want to test spatial hypotheses about aging and AD, you need data that preserve the cortex as a spatial object. This method is a step toward making such data available.

## Methods

### Animals

Two adult female rhesus monkeys (*Macaca mulatta;* A27, 22 years; A26, 24 years) from the Oregon National Primate Research Center were used in this study. Both were pre/peri-menopause at the time of tissue harvest; for hormone assay data, health history, and prior research participation, see **Supplementary Information**. After moving to Duke University, animals were housed with grooming bar access. Diurnal cycling was supported by a 12:12 light cycle. Animals were fed LabDiet 5038 biscuits and fresh fruits and vegetables in-cage, in addition to small treats (nuts, dried fruit, yoghurt drops, etc.) used as reinforcers for cognitive testing. Water was provided *ad libitum*. Neither food nor water access restriction was employed as part of behavioral testing. Animals were Simian B-virus, tuberculosis, and measles negative, and were maintained at body condition scores of 3.0-4.0. Experimental procedures described below were conducted under a protocol approved by the Duke Institutional Animal Care and Use Committee (IACUC) and followed guidelines published by the National Institutes of Health.

### A26/A27 premortem studies

Prior to tissue harvest, animals completed cognitive assays (delayed recognition span and delayed response tasks), 3T magnetic resonance imaging, and collection of plasma and cerebrospinal fluid (CSF) samples as part of a larger study not described here.

### Tissue sample collection

Animals were initially sedated with ketamine (20 mg/kg, IM), prepared for surgery (including intubation and placement of an IV catheter), and placed in a stereotaxic frame. Final premortem blood and CSF samples were collected by veterinary staff, and then anesthesia was induced and maintained with inhaled isoflurane (1-2.5%) and mechanical ventilation. To facilitate rapid harvest, a craniotomy was prepared to expose the cortex bilaterally from just posterior to the coronal suture to the occipital ridge. A small bridge of bone was left intact spanning the midline anteriorly and posteriorly, and the bony plate left in place.

Once the craniotomy was complete, the animal was removed from the stereotaxic frame, maintaining ventilation, and euthanized by propofol overdose (10 mg/kg IV) prior to rapid (> 300 mL/min) transcardial perfusion with chilled 0.1 M phosphate-buffered saline (PBS, pH 7.42 in type 1 water). 1000 IU heparin was added to the first liter of PBS to prevent clotting and facilitate clearing of the vasculature. When the solution exiting the right atrium ran clear, and the tongue and liver were visibly exsanguinated, the remaining bony bridges across the midline of the skull were clipped, the dura opened, and the brain removed. The brain was bisected along the midline.

One (randomized) hemisphere was immersion fixed for unrelated anatomical studies. The cortex of the other hemisphere (A26, right; A27, left) was separated from subcortical structures, cooled over dry ice, and approximately flattened (up to the end of step 7 in the protocol described by Sincich et al.^19^). The flattened cortex was then voxelated by hand, on dry ice, into small (A26: ∼4 mm/side; A27: ∼2.5mm/side) tissue blocks. To counterbalance order of freezing effects, A26 was dissected beginning at the frontal pole, and A27 at the occipital pole. Each sample was immediately placed in a cryotube and stored on dry ice. After collection was completed, samples were stored at -80°C. The first sample/last sample delays (hh:mm) from anoxia to freezing were: A26 = 00:41/02:23; A27 = 00:42/02:34). Video footage of the dissection was later used to reconstruct each sample’s position in the cortical sheet (see **Fig. 1**).

**Figure 1:**
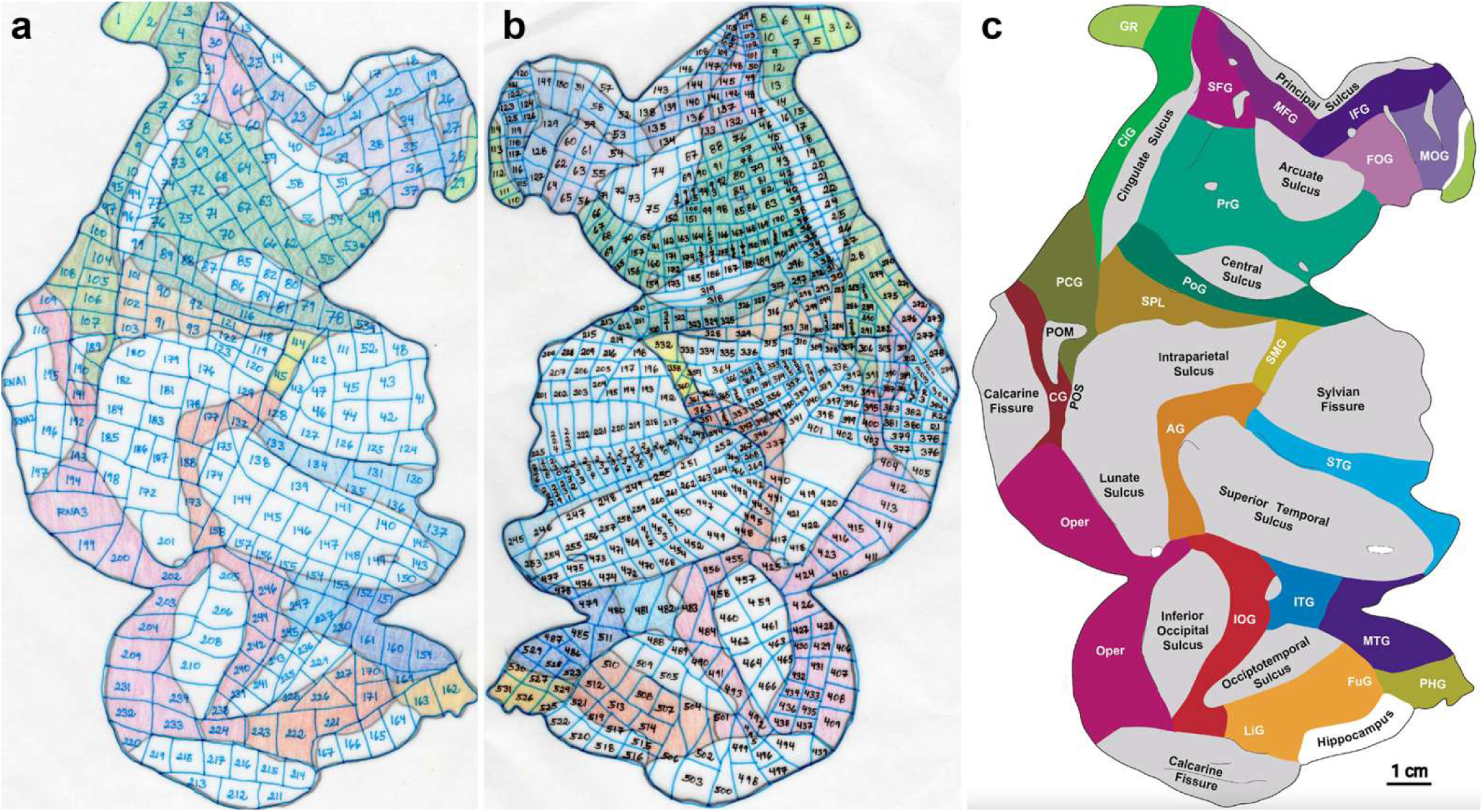
Cortical sampling. **(a)** Flatmap for A26 (right hemisphere). **(b)** Flatmap for A27 (left hemisphere, map reversed). Hand drawn maps in panels (a) and (b) manually colored to roughly match the scheme in panel (c). **(c)** Flatmap outline and brain regions from Figure 4 in Sincich et al.^19^. Gyri in color and sulci in gray; reproduced with permission. Gyri: GR, gyrus rectus; CiG, cingulate gyrus; SFG, superior frontal gyrus; MFG, middle frontal gyrus; IFG, inferior frontal gyrus; FOG, fronto-orbito gyrus; MOG, medial orbital gyrus; PrG, precentral gyrus; PoG, postcentral gyrus; PCG, pre-cuneate gyrus; SPL, superior parietal lobule; SMG, supra-marginal gyrus; CG, cuneate gyrus; AG, angular gyrus; STG, superior temporal gyrus; Oper, operculum; IOG, inferior occipital gyrus; ITG, inferior temporal gyrus; MTG, middle temporal gyrus; PHG, parahippocampal gyrus; LiG, lingual gyrus; FuG, fusiform gyrus. Sulci: POM, parieto-occipital sulcus (medial); POS, parieto-occipital sulcus.

### Exsanguination control

Data from a third animal was included to assess the impact of exsanguination on metabolomic profiles. This juvenile (∼2.5 years) male rhesus monkey (U5) had undergone electrophysiological brain recording under an IACUC-approved protocol at Johns Hopkins University^20^. Following euthanasia, the brain was harvested without exsanguination (i.e., no transcardial perfusion), flash-frozen using isopentane (approximate PMI ∼10 minutes), and stored at −80°C. The frozen brain was transported to Duke University on dry ice and stored at −80°C until sampling (∼5 months post-harvest). On the day of sampling, the intact (non-flattened) left (unrecorded) hemisphere was sampled anterior-to-posterior by 3-mm punch biopsy, through a grid applied to the surface of cortex (still frozen), yielding 75 tissue punches (see **Supplementary Fig. S1**). Tissue samples from U5 were profiled using the same Biocrates Q500 kit and processing pipeline as described for A26 and A27 below except that during homogenization, samples were extracted at a 1:4.5 ratio instead of 1:10. Appropriate corrections were made in the analyses.

### Tissue sample preparation

Samples were weighed in their cryotubes and an estimated weight calculated based on an average tube weight of 1.1354 g. The median initial sample weight was 130 mg. Samples with an estimated weight < 50 mg were combined with their lowest-weight neighbor. Samples > 120 mg were split into two, and those > 240 mg into three. Split samples were treated as held-out data/replicates for their unique mapped location (see **Fig. 1** for maps), not assigned to a new location. Sample order was randomized (random.org) and blinded at this point.

Brain tissue was homogenized in CK14 tubes (Bertin) with an additional 50-mm stainless-steel ball (Qiagen) and extracted by adding methanol:water:chloroform (67:15:18) at a 10:1 ratio (v/w), followed by homogenization with at least two cycles (2 × 30 s at 30 Hz) using a TissueLyserIII (Qiagen) and centrifugation (10,000 × g, 3 min, 4°C); supernatants were transferred to 2-mL Protein Lobind plates (Eppendorf) and stored at −80°C prior to analysis.

### Study pool quality control sample generation

A study pool QC (SPQC) was generated by pooling aliquots across study samples. SPQC samples were interspersed across plates as measures of total analytical variability within and across plates/batches. In addition, for proteomics, a post-digestion pooled QC was generated by pooling digests (A26: plate-by-plate; A27: from plates 1 & 2 only) to assess post-digestion technical reproducibility and plate-level performance.

### Targeted metabolomics

Samples (A26: n = 406; A27: n = 549) were profiled using the Quant 500 targeted metabolomics kit (Q500; Biocrates). Q500 plates were prepared in accordance with the manufacturer’s protocol: internal standards were pre-applied to each well; 10 µL each of blanks, calibration standards, Biocrates QC materials, and a global reference QC were added to designated wells; and 30 µL of each sample supernatant loaded. Plates were dried under nitrogen (30 min), derivatized with phenyl isothiocyanate, and eluted with 5 mM ammonium acetate in methanol. For ultra-high-pressure liquid chromatography (UHPLC) analysis, eluates were diluted 1:1 in 80:20 water:methanol; for flow injection analysis (FIA), eluates were diluted 20:1 in the kit running solvent. Each plate included three levels of Biocrates-provided quantitative human plasma QC materials, a pooled study QC (SPQC), and a reference plasma (GoldenWest). The SPQC pool was generated and injected once before, once during, and once after the study samples. SPQC measurements were acquired in replicate across LC and FIA modes.

UHPLC separations were performed on a Waters Acquity UPLC I-Class using a 2.1 × 50 mm, 1.7 µm BEH C18 column (with guard) using a gradient from 0.2% formic acid in water to 0.2% formic acid in acetonitrile (≈6 min/sample). Two UHPLC injections were acquired (ESI+ and ESI−) using MRM on a Sciex TQ/S 6500+ platform. FIA-MS/MS measurements (e.g., lipids, acylcarnitines, and neutral lipid classes) were acquired as separate FIA methods (≈3.8 min/injection) in ESI+ on a Sciex QTRAP 6500+ with sample introduction via the Acquity system. Raw concentration outputs (pmol/mg tissue) were exported using the Biocrates software platform which provided peak integration, signal evaluation, calibration curve building, and result validation. Analyte-level status flags (e.g., valid, <LOD, internal standard out of range, no peak detected) were retained to support downstream filtering and sensitivity analyses.

### Metabolomics quality controls (QC) and batch correction for cross animal study

Each animal was analyzed separately yielding two analytic batches. Within each of these, samples spanned multiple plates. Across both analytical batches, performance was monitored using Biocrates QC Level 1–3 materials, a pooled study QC, and a reference plasma (GoldenWest), run repeatedly across plates to track preparation and instrument variability. In exploratory QC analysis, PCA performed within compound classes showed SPQC injections clustering centrally relative to study samples. SPQC samples were used for within-animal batch correction; between-animal batch correction was based on subtraction of the Level 3 Biocrates QC materials.

### Untargeted proteomics

Protein concentrations were determined by detergent-compatible Bradford assay (Pierce) and extracts normalized (A26: 0.6 µg/µL; A27: 0.4 µg/µL) using an Opentrons OT-2 liquid-handling workflow. Samples were reduced (10 mM dithiothreitol, 80°C, 15 min), alkylated (25 mM iodoacetamide, room temperature, dark), and 10 µgs of protein was processed using S-Trap 96-well plates (Protifi) or S-trap micro devices using the manufacturer’s protocol. The digestion step (1.5 h at 47°C) used 1 µg modified bovine trypsin (Worthington Sequenz) in 25 µL triethlammonium bicarbonate, pH 8.5 (TEAB). Peptides were eluted (40 µL TEAB, then 2 x 40 µL of 0.2% formic acid), and yeast alcohol dehydrogenase (ADH; 50 fmol final) was included as a QC spike for downstream monitoring. Ten µL (approximately 600 ng) of individual samples and SPQC replicates were loaded onto disposable Evotip Pure trap columns for analysis.

Peptides were analyzed using an Evosep One LC with an 8 cm × 150 µm PepSep column (1.5 µm particles) and PepSep sprayer/emitter, interfaced to an Orbitrap Astral MS (ThermoFisher). MS1 scans were acquired at 240,000 resolution over 380–980 m/z with automatic gain control (AGC) target 500% and 50 ms maximum injection time (IT); data-independent acquisition (DIA) MS/MS used 4 m/z isolation windows over the same precursor range with AGC target 500%, 6 ms IT, and normalized collision energy of 28% (default charge state 3; RF lens 40%). Raw files were converted to *.htrms and processed in Spectronaut 19 using direct-DIA to build study-specific spectral libraries based on a UniProt *Macaca mulatta* database (RRID:SCR_002380, taxon ID 9544; downloaded 5/25/24) appended with common contaminants (via FragPipe). A single spectral library was built from combined search archives. Searches used trypsin/P specificity (≤ 2 missed cleavages; peptide length 7–52 aa) with fixed carbamidomethyl (Cys). Identifications were filtered at 1% precursor and protein group FDR (q-value). Protein abundances were calculated using MaxLFQ and normalized to the global median in a single analysis batch using Spectronaut.

### Cortical sheet reconstruction and conversion to matrix

Reconstructed maps (see *Tissue sample collection* and **Fig. 1**) were overlaid on graph paper and sample boundaries re-drawn with right angles within a matrix containing 100 x 60 (height x width) square cells.

### Selection of samples for statistical analysis

Samples included in the final analysis were selected through a multi-step, data-driven procedure designed to ensure quality, consistency across modalities, and robustness to outliers. For both A26 and A27 datasets, metabolomic and proteomic data were first preprocessed and batch-corrected independently. Quality control procedures included filtering of metabolites with excessive missingness (threshold: missing from 70% of samples for triglycerides, 85% for all others), imputation using half the limit of detection (LOD/2), adjustment for dilution factors, and log-transformation. Batch effects were corrected using ComBat. In metabolomics, additional correction using SPQC samples was applied prior to batch correction by subtracting the SPQC (plate-wise). SPQC adjustment was not applied for proteomics, based on empirical assessment (see **Supplementary Fig. S2**).

To construct a unified set of samples across both modalities, we implemented a held-out preprocessing and sample selection procedure. When two samples were available for a single map position (due to splitting, see above), we retained the one more consistent with the overall data distribution. Specifically, we quantified the extremeness of each sample using a robust PCA-based distance metric. For each dataset (metabolomics and proteomics), we computed the first two principal components (PC1 and PC2) and defined the distance of a sample from the robust center as:

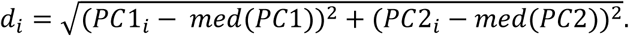

This distance was then normalized using the median absolute deviation (MAD):

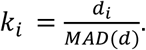

The statistic *k*_i_ measures how far a sample lies from the central structure of the data and is then taken the maximum between the metabolomics and proteomics. When multiple candidate samples were available, we retained the sample with the smaller *k*_i_ value, thereby favoring samples that are more representative of the global data structure.

Finally, we included only samples for which both metabolomic and proteomic data was available. This resulted in 233 matched samples for A26 and 508 matched samples for A27.

### Feature clustering guided by principal components

Both the proteomic and metabolomic assays yield high-dimensional data with strong collinearity among features. Many variables exhibit substantial correlations with each other, making it challenging to distinguish their contributions in downstream analyses. To address this issue, we developed **PChclust**, a feature-clustering algorithm guided by principal components (PCs) that aggregates co-linear variables into representative clusters.

We applied PChclust separately to each modality (metabolome and proteome) using a consistent framework, enabling downstream integration at the level of cluster-derived features. Within each dataset, all variables were first centered and standardized. We then performed hierarchical clustering based on pairwise similarities between variables (Spearman correlation). The resulting dendrogram serves as the initialization. Starting from the root node of the dendrogram, for each potential split, we conducted eigendecomposition on the set of variables before the split. The split was accepted only if the proportion of variance explained by the first PC was below a predefined threshold of 70%. Otherwise, if the top PC already explained more than 70% of the variance, we stopped splitting that branch. This procedure was applied recursively until either the cutoff criterion was met or only single variables remained at the leaf nodes.

Unlike standard hierarchical clustering, which relies on cutting the dendrogram at a fixed depth, our approach adaptively stops splitting when the leading PC sufficiently summarizes the variance structure of a variable set. For two standardized variables with correlation *ρ*, the variance explained by their first PC is (1 + *ρ*)/2. Thus, PChclust naturally limits inter-cluster correlation and ensures that the resulting clusters are relatively independent. The output of PChclust is a collection of variable clusters, each represented by its first principal component, along with singlet variables that are not strongly correlated with any other variables or clusters.

### Spatial canonical correlation analysis

Canonical correlation analysis (CCA) studies the relationship between two sets of variables by finding linear combinations of each set that maximize their correlation. Traditional CCA typically assumes that samples are independent. However, in our data, samples exhibit spatial dependencies that must be taken into account. To address this, we developed **spatially regularized sparse CCA (sr-sCCA)**, a method that incorporates spatial information into the extraction of canonical components to enhance their spatial relevance.

Let *X* ∈ *R*^n×p^ and *Y* ∈ *R*^n×q^ denote the metabolomic and proteomic data matrices of *p* and *q* features measured across *n* spatially distributed samples. We first constructed a spatial map of the *n* samples based on the maps of sample position on the cortical sheet (see *Cortical sheet reconstruction and conversion to matrix* and **Fig. 1** for the raw maps). Using Delaunay triangulation on the samples’ x-y coordinates, we define a spatial neighborhood matrix *W*_0_ ∈ *R*^n×n^, where (*W*_0_)_ij_ = 1 if samples *i* and *j* are spatial neighbors and 0 otherwise. Let *D* be the diagonal degree matrix of *W*_0_, and define the normalized graph Laplacian as:

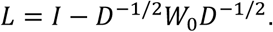

Denote *L* = *Q*Ʌ*Q*^T^as the eigendecomposition of *L*. We then construct a spatial smoothing matrix:

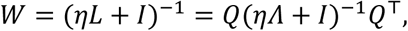

where 5 = 1/*median*(*diag*(Ʌ)). The matrix *W* is positive definite and serves to encode spatial smoothness among samples.

Borrowing ideas from spatial PCA, our sr-sCCA then seeks sparse canonical vectors *a* ∈ *R*^p^ and *b* ∈ *R*^q^ that maximize the following objective function:

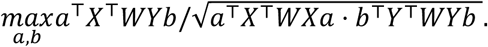

Because *W* is positive definite, this optimization can be implemented using existing sparse CCA solvers by applying a spatial transformation to the data

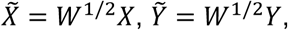

and then performing standard sparse CCA on 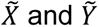. This transformation enables joint analysis of metabolomic and proteomic data and ensures that spatial relationships among samples are incorporated into the estimation of canonical components, leading to spatially coherent and interpretable results.

### Cross-animal component projection

The A27 canonical loading vectors (10 components) were mapped back from cluster space to the original feature space and then feature-level weights restricted to features common between animals (9,250 proteins; 277 metabolites). These weights were then applied to the A26 proteomic and metabolomic data matrices. The resulting scores were projected onto the A26 cortical spatial map (**Fig. 1a**) to generate transferred spatial score maps for each component.

### Analysis of sampling resolution

To assess spatial similarity in molecular profiles—and thus determine adequate sampling resolution for this method, based on the different sampling densities of the two animals (**Fig. 1**)—we computed correlations between each sample and its spatial neighbors for both the metabolomic and proteomic datasets. First, spatial adjacency relationships among samples were constructed based on the Delaunay triangulation (see above). This adjacency structure defines first-order neighbors as directly connected samples. Higher-order neighbors were derived algebraically from the adjacency matrix: second-order neighbors correspond to samples connected through two-step paths, and third-order neighbors correspond to three-step paths while excluding lower-order connections, and so on.

Metabolomic and proteomic feature matrices were first restricted to samples present in the spatial network and then standardized by mean-centering and scaling each feature to unit variance. For each pair of samples belonging to a given neighbor order, we computed the Spearman correlation between their molecular profiles using pairwise-complete observations. The correlations with neighbors of the same order were averaged for each sample and visualized on the brain map (**Fig 4**).

### Analysis for exsanguination control study

To assess the effect on metabolomic profiles of the presence of blood in the samples, we compared metabolomic data from A26 and A27 (perfused, exsanguinated) with U5 (blood present). Each animal’s data was independently preprocessed using the pipeline described above. To enable cross-animal comparison, data were normalized by subtracting per-plate means of the Biocrates QC Level 3 materials—a standardized reference common across all kits. The intersection of metabolites passing QC filters in all three animals yielded 252 common metabolites. Formal batch correction (ComBat^21^) was not applied to the combined dataset; ComBat assumes samples are drawn from a common distribution and thus would remove the biologically meaningful between-animal variance that was the target of this analysis. PCA was performed on the combined, QC3-normalized data (**Fig. 5**). The PC driving the separation between perfused and unperfused tissue (PC1) was then correlated with the component loadings from the spatial CCA analysis using Spearman correlation coefficients across the set of shared metabolites.

### Biological pathway enrichment analysis

In order to understand the biological significance and aggregate molecular signatures of the proteins in each cluster, we employed the Metascape program (metascape.org)^22^. The genes associated with each protein cluster were input as the target gene list. In order to group the top 20 enriched Metascape output GO terms into broader biological categories, functions of each individual gene (based on literature search) contributing to a particular category were qualitatively grouped into major functional categories. If a majority of the target genes associated with an enriched pathway GO term were associated with a particular functional category, this category was assigned to the GO term. A single GO term could be assigned to multiple functional categories. Minimum overlap of genes was 3, and a p-value cutoff of 0.01 and minimum enrichment of 1.5 were used. For pathway databases, the GO biological processes, reactome gene sets, KEGG pathways, and WikiPathways were used; for structural complexes, the CORUM database was used. Manual annotation was used to assign a label (e.g., synaptic processes, mitochondrial processes) to each analysis result.

### Integrated proteomics and metabolomics analysis

The strategy for integrated analysis of the proteomic and metabolomic data from the samples was to utilize a “proteomics-driven” approach. The list of proteins from three PChclust clusters were used in example joint pathway analyses; one focused on each of three biological pathways identified by the above enrichment analysis (see Results for selected pathways). The average metabolite abundance was used for the list of targeted metabolites in the joint analysis. The Joint Pathway Analysis implemented in MetaboAnalyst^23^ is based on weighted integration. The weight strategy for different omic domains (“proteomics data universe” and “metabolomics data universe”) is performed with a weighted z-test. The weighted z-test is proposed for combination analysis and leveraged for the weighted integration of different datasets with significantly different sizes. Weights are assigned based on the proportion of proteins and metabolites in the specific domain to balance the influence from the different sizes of the input lists. The adjusted p-values for the joint analysis are estimated from the weighted z-test.

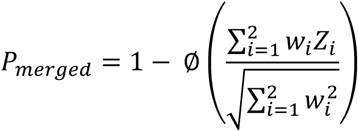

*w*_i_ are the weights of the *p-*values of proteins or compounds within individual omics universe or pathway domains (i = 1 for proteomics, 2 for metabolomics), respectively; Z_i_ is the Z score of the corresponding p-values of single omics (protein or metabolite) data, Z_i_ = Φ^−1^(1 −P_i_); Pi are the p-values from the enrichment analysis; Φ denotes the standard normal cumulative distribution function.

The pathway impact score is a graph theory-based measure that represents a combination of the centrality of the pathway in the network and the pathway enrichment result; these are calculated as an importance measure. The pathway impact score is calculated as the sum of the importance measures for the matched proteins/metabolites in the network normalized by the sum of importance measures of all proteins/metabolites in the pathway. The pathway impact score represents an objective estimate of the importance of a given pathway relative to a global metabolic network.

In addition to the proteomic-driven joint pathway analysis, we also performed an integrated joint pathway analysis using all proteins and metabolites. Loadings for the first PC for both proteins and metabolites were used as the scalar values (proxy for fold change) for the analysis.

### Biomarker assays on CSF samples

CSF samples from A26 and A27 (plus ∼20 additional rhesus monkeys) were profiled using the NULISAseq CNS Disease Panel 120 (Alamar Biosciences, Fremont, CA), a 120-plex targeted immunoassay based on the NULISA (NUcleic acid Linked Immuno-Sandwich Assay) platform^24^.

Samples (25 µL CSF input) were stored at −80°C until the assay, then thawed and centrifuged at 2,200 × g for 10 min. All 45 macaque samples were run on a single plate on the ARGO HT system, with library sequencing on the Element Biosciences AVITI platform. Data were normalized using internal and inter-plate controls and are reported as NULISA Protein Quantification (NPQ) units (log2-transformed normalized counts). Per-animal mean NPQ values for A26 and A27 were compared to the full macaque cohort by z-score to screen for outlier biomarker profiles.

## Results

The entire cortical sheet of one brain hemisphere was flattened and voxelated by hand-dissection for two animals, yielding 237 (A26) and 511 (A27) unique mapped cortex locations (**Fig. 1**). Each tissue voxel was homogenized and split to yield matched samples for targeted metabolomics and an untargeted proteome. After excluding outliers and lost samples, we had 233 (A26) and 508 (A27) samples with matched metabolomic and proteomic data.

### Biomarker analysis

Cerebrospinal fluid (CSF) samples (A26 = 2 samples; A27 = 3), were submitted to Alamar Biosciences for NULISA^24^ analysis as part of a larger study (reported separately). Briefly: Of 129 panel targets, 79 (61.2%) were detectable in ≥ 50% (of 45 total, ∼20 individuals) macaque samples, all run on a single plate. To assess the neurological and inflammatory status of A26 and A27, mean NULISA Protein Quantification (NPQ) values for each animal were compared to the full cohort by computing z-scores for each detectable target. Both animals fell well within the population distribution across nearly all targets. A26 had a single target (IL-7) with *z* > 2 (*z* = +2.9; ∼2.2-fold elevation). A27 had no targets exceeding *z* > 2. Key neurodegenerative biomarkers for both animals—GFAP, neurofilament light and heavy chain, total tau, phospho-tau species, NPTX2, CHI3L1 (YKL-40), and alpha-synuclein—were all within one standard deviation of the population mean. Thus, neither animal exhibited evidence of neurodegeneration, neuroinflammation, or CNS immune activation at the time of CSF collection and tissue harvest.

### Targeted metabolomics: analytical performance and quality control

Targeted metabolomics was performed using the Biocrates MxP Quant 500 (Q500) kit. Each animal was analyzed independently across multiple plates (A26: 6 plates, A27: 7 plates; see **Table 1**), each of which included human plasma quality control materials and a study pool quality control sample (SPQC, see Methods).

**Table 1.**
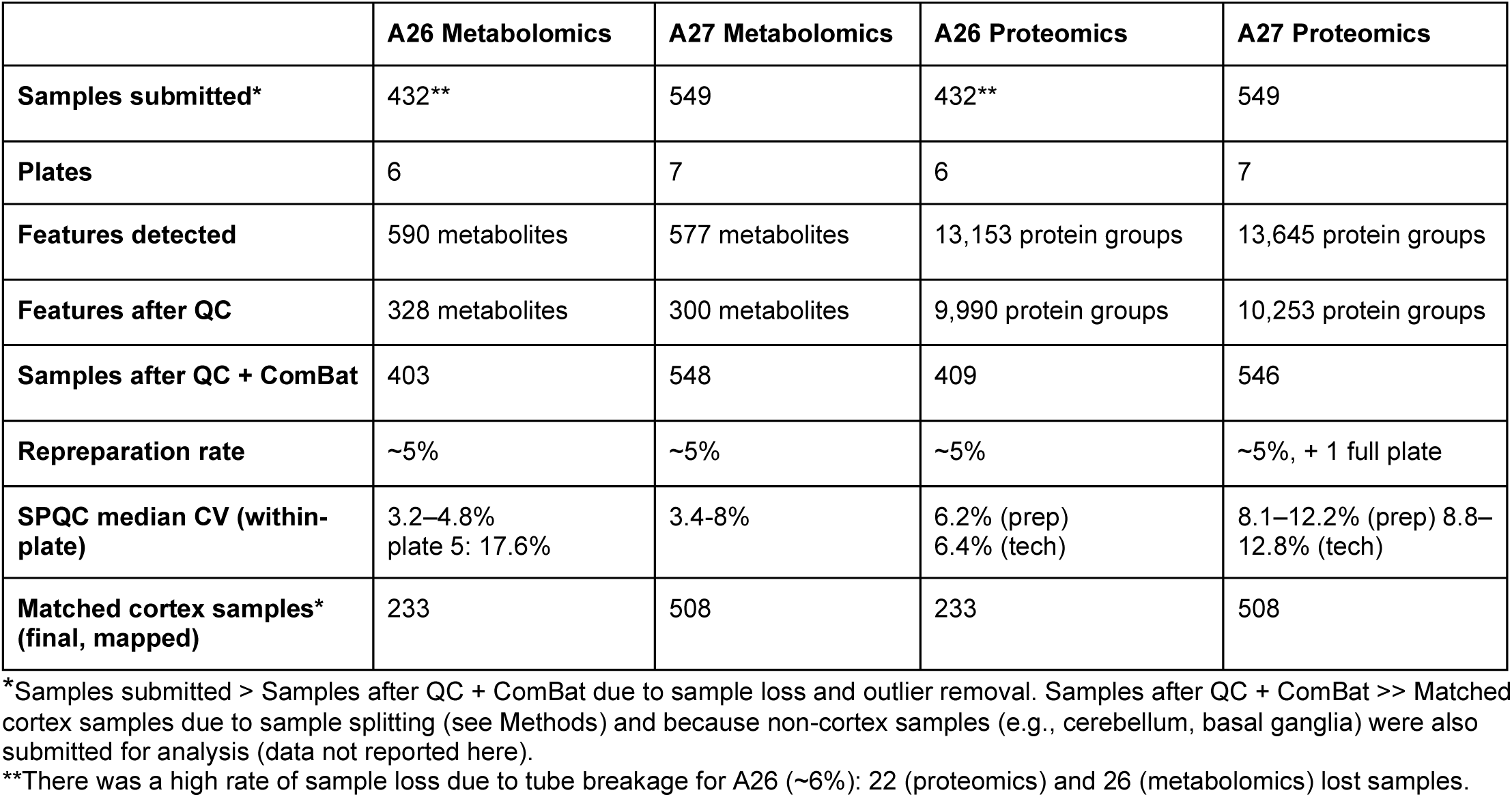
Summary of LC-MS/MS analytical performance.

#### Analyte screening

Of 630 targeted metabolites, 590 (A26) and 577 (A27) were detectable in the study samples. After removing analytes with excessive missingness (below LOD in > 70% of samples for triglycerides; > 85% for all other classes), 328 analytes were retained for A26 and 300 for A27.

#### Sample screening

Outlier samples (A26 = 3; A27 = 1) were removed before batch correction by ComBat^21^. A further 4 samples from A26 were removed after batch correction. Within-plate technical reproducibility was high (see **Table 1**). Triglycerides and fatty acids showed higher variability than other classes (17–21%), consistent with the lower precision typically observed for FIA-measured lipid classes^25^ and the low native abundance of triglycerides in brain parenchyma. Principal component analysis (PCA) confirmed central clustering of SPQC replicates relative to study samples for both animals (**Supplementary Fig. S2a,c**).

### Untargeted proteomics: analytical performance and quality control

Untargeted data-independent acquisition (DIA) proteomics was performed after chromatographic separation. Each animal was processed across multiple plates (A26: 6 plates; A27: 7 plates; **Table 1**), with study pool QC replicates (SPQC_prep) interspersed at 4 per plate; post-digestion pooled QC replicates (SPQC_tech); and a 50 fmol yeast ADH spike in each sample.

#### Analyte screening

Spectral library construction in Spectronaut 19 yielded 13,153 protein groups for A26 and 13,645 for A27. After quality filtering, 9,990 protein groups were quantified for A26, and 10,253 for A27. Per-sample identification rates averaged ∼8,250 protein groups per plate for A26 and ∼8,600 for A27.

#### Sample screening

Approximately 5% of samples in each animal required repreparation due to incomplete digestion or failure to meet analytic criteria (e.g., low identifications); for A27, one plate (plate 2) was reprepared in its entirety due to potential detergent contamination. Technical reproducibility was excellent: for example, in A26, median CVs were 6.2% (SPQC_prep) and 6.4% (SPQC_tech), compared to 21.6% for individual study samples. PCA showed tight clustering of SPQC replicates with some plate-driven separation for both animals (**Fig. 2b,d**). Data were normalized and protein abundances calculated using MaxLFQ. Cross-animal comparability was supported by inclusion of A26 SPQC material alongside A27 study samples.

**Figure 2:**
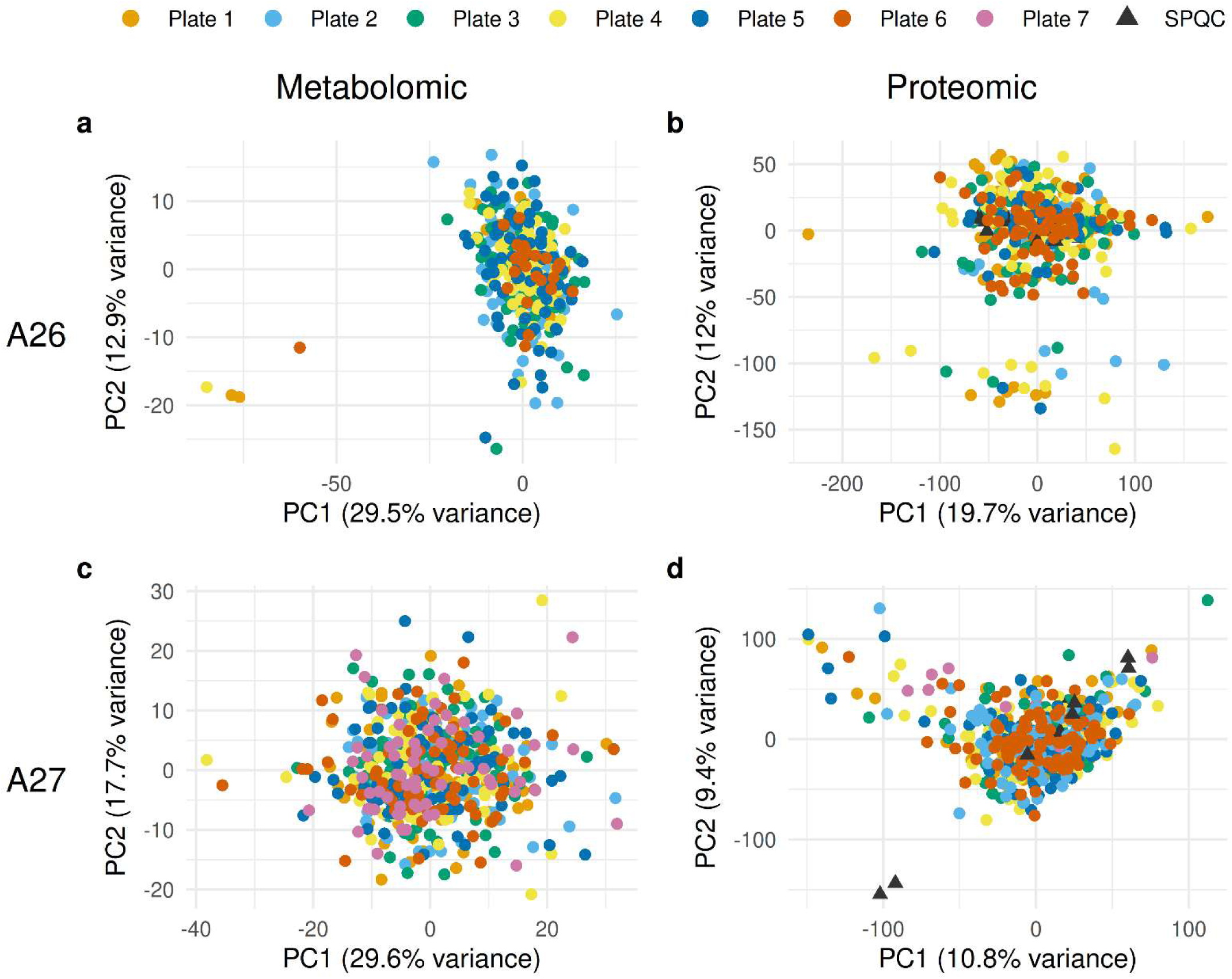
Principal component analysis (PCA) confirms removal of batch effects by ComBat^21^. PCA of batch-corrected data for metabolomics **(a,c)** and proteomics (**b, d**) in A26 (top row) and A27 (bottom row). Each point represents a single sample, colored by processing plate. Black triangles (**b, d**) indicate study pool quality control (SPQC) replicates. Central clustering of SPQC replicates relative to the spread of study samples (b, d; for metabolomics see **Supplementary Fig. S2a,c**) confirms that technical variability is small compared to biological variability between samples.

### Spatially regularized sparse canonical correlation (sr-sCCA) analysis

Because neighboring cortical voxels are not statistically independent, standard multivariate methods that assume independent samples are not appropriate for our data. We developed a spatially regularized sparse canonical correlation analysis (sr-sCCA, see Methods) that incorporates spatial neighborhood structure via graph Laplacian smoothing, penalizing solutions in which nearby voxels have very different canonical scores. **Figure 3** shows the spatial distributions of the first (V1, U1) and second (V2, U2) pairs of canonical variates derived from the sr-sCCA analysis of the metabolomic and proteomic profiles in A26 and A27 (**Supplementary Fig. S3-4** show all 10 components). In both animals, the metabolomic and proteomic canonical scores exhibit broadly similar spatial patterns: a mediolateral gradient for the first V and U components (**Fig. 3a,c,e**), and an anteromedial-posterolateral gradient for the second (**Fig. 3b,d,f**). Regions with higher metabolomic scores generally corresponded to regions with higher proteomic scores, suggesting that the dominant shared signal between the metabolome and proteome is spatially structured across the cortical surface.

**Figure 3:**
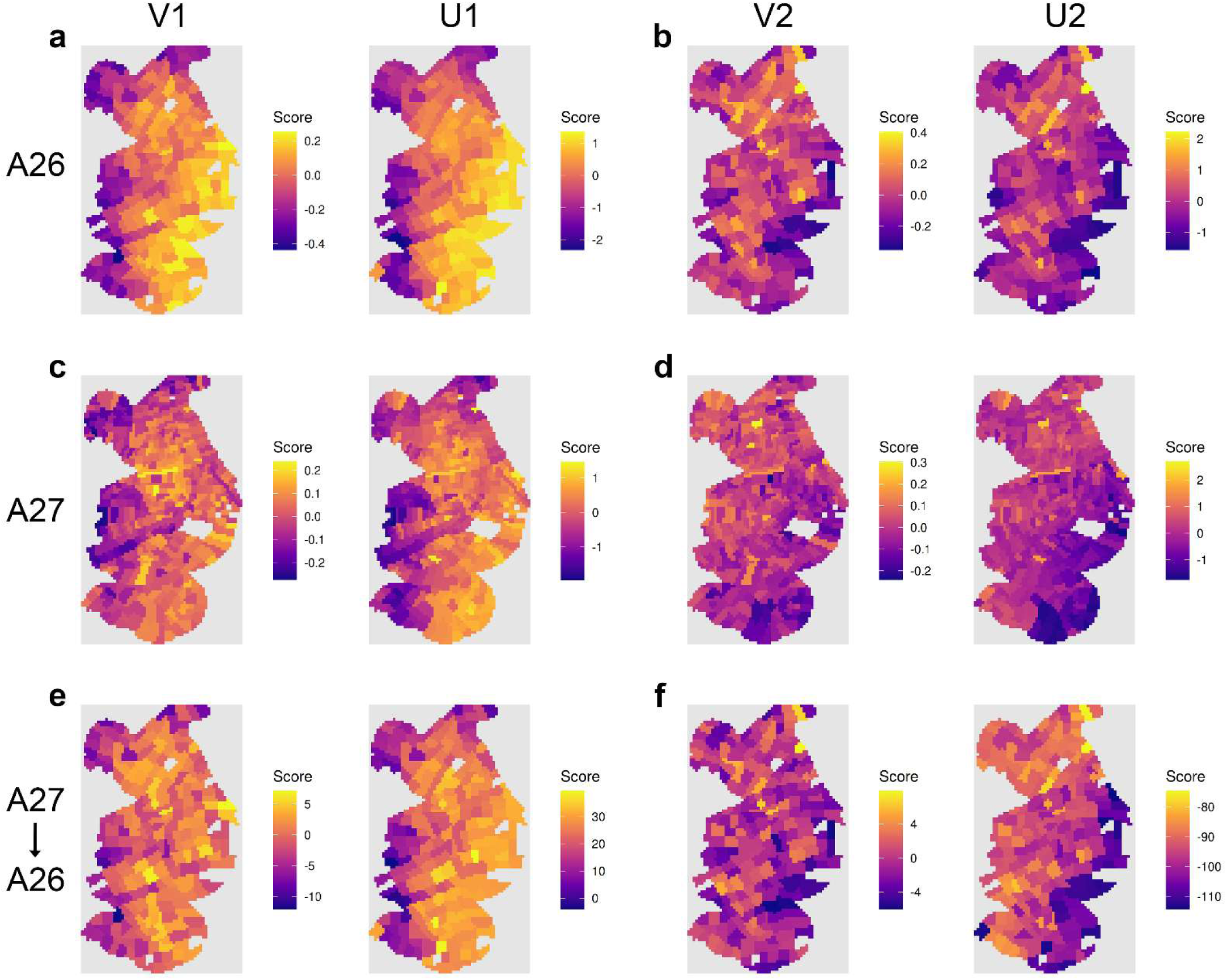
Spatial correlation gradients. Spatial maps created by projecting metabolomic canonical scores (V) and proteomic canonical scores (U) from spatially regularized sparse canonical correlation analysis (sr-sCCA) onto the cortical spatial coordinates for each sampled location. Note that color scales are independently normalized for each component and represent relative component scores within each map; as such, colors are not directly comparable across maps. Warm colors (orange-yellow): high/positive correlation scores; cool colors (pink-purple): low/negative correlation scores. **(a)** Spatial maps for the first component’s metabolomic (V1) and proteomic (U1) scores in A26, illustrating coherent spatial variation across modalities. **(b)** As in panel A, for the second component (V2/U2). **(c)** As in panel A, for A27. **(d)** As in panel B, for A27. **(e)** As in panel A, but with A27-derived sr-sCCA canonical vectors for component 1 (V1/U1) projected onto A26 data. **(f)** As in panel E, for component 2 (V2/U2).

### Spatial decay of molecular similarity: sampling density study

The two animals were voxelated at different resolutions (∼4 mm for A26, ∼2.5 mm for A27; see **Fig. 1**) in order to assess the sampling density required to capture spatial molecular organization. The distributions of average correlations across samples (see Methods; **Supplementary Fig. S5**) are summarized in **Figure 4**. Across both metabolomic and proteomic data, the average correlation decreases as neighbor order increases, indicating a spatial decay of molecular similarity.

**Figure 4:**
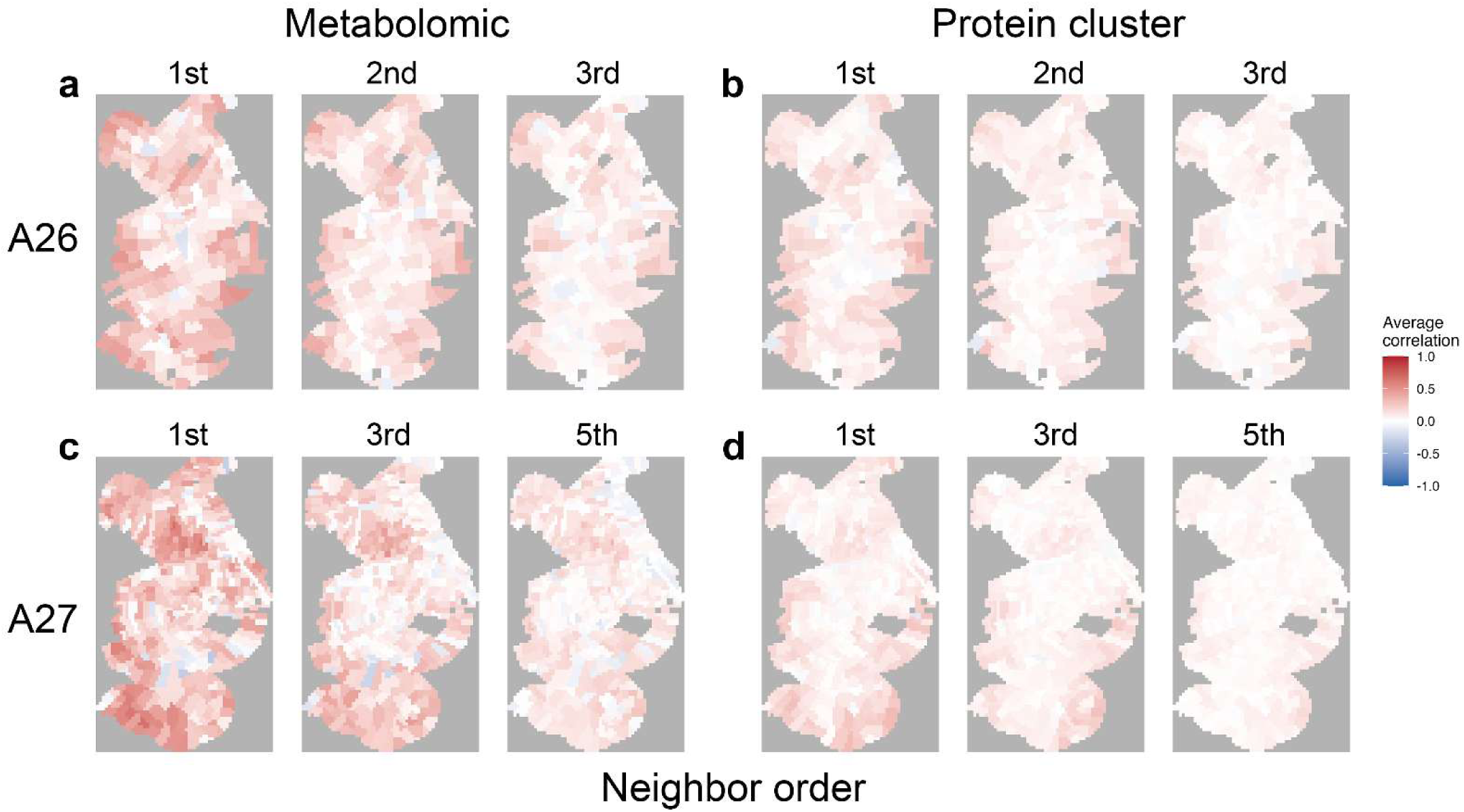
Spatial decay of correlations with neighboring samples. Each panel shows a spatial correlation map in which each sample is colored by its average Spearman correlation with neighbors at the indicated neighbor order (1st, 2nd, 3rd for A26; 1st, 3rd, 5th for A27). Warmer colors indicate higher local similarity; cooler colors indicate greater local heterogeneity. Neighbor orders are defined by steps through the Delaunay triangulation of sample positions (see Methods). **(a)** Metabolomics, A26. **(b)** Proteomics, A26. **(c)** Metabolomics, A27. **(d)** Proteomics, A27. Both sampling resolutions captured spatial decay of molecular similarity, but A27’s denser sampling (∼2.5 mm) resolved regional heterogeneity in the metabolomic data that was not detectable at A26’s coarser (∼4 mm) resolution. This difference was less pronounced for proteomics.

The spatial correlation maps in **Figure 4** display each sample’s average correlation with its neighbors. As the neighbor order (i.e., physical distance) increases, the spatial patterns become more diffuse, reflecting the reduced similarity among more distant samples. Both sampling resolutions detected spatial decay of molecular similarity (**Fig. 4**), but not equivalently. In the metabolomic data, A27’s denser sampling resolved regional heterogeneity in correlation structure (**Fig. 4c**, left) that was not detectable in A26. Correlation maps at A27’s 3rd-neighbor order (**Fig. 4c**, middle), which correspond roughly to A26’s 1st-neighbor physical distance (**Fig. 4a**, left), still showed spatially differentiated warm and cold patches. A26’s correlation maps were largely uniform by 2nd-neighbor order. This difference was less pronounced in the proteomic data, where both animals showed comparable spatial decay profiles (**Fig. 4b,d**).

### Cross-animal reproducibility of spatial canonical components

The sr-sCCA analysis identifies canonical components that capture spatially coherent variance, but each animal’s analysis is performed independently, so the resulting maps could reflect idiosyncratic features of the individual animal, rather than conserved biology. To test this, we projected the A27-derived sr-sCCA canonical vectors onto A26 data and asked whether coherent spatial maps emerged (**Fig. 3e,f**).

For both components, projecting the A27-derived vectors onto A26 recovered the spatial pattern seen in A26’s own sr-sCCA (compare U1 and U2 in **Fig. 3a,b** with **3e,f**); for component 2, the projected proteomic map showed the anteromedial-posterolateral gradient more clearly than A26’s own fit. The projected metabolomic scores were spatially noisier than the proteomic scores. Because the A27 vectors were estimated without reference to A26, this agreement indicates that the dominant shared spatial structure of the two molecular layers is conserved across the two animals.

### Effect of exsanguination on metabolomic profiles

Human brain samples are not exsanguinated during harvest. To understand the extent to which an analysis of samples containing blood would differ from that presented above, we compared the metabolomic profiles for A26 and A27 with that of a third animal (U5) whose brain was harvested without perfusion. After independent preprocessing and cross-animal normalization using Biocrates QC standards (see Methods), 252 metabolites common to all three animals were retained for comparison.

PCA of the combined, QC3-normalized data showed clear separation of U5 from A26 and A27 along PC1, which accounted for 40.2% of the total variance (**Fig. 5**). A26 and A27 samples were intermixed along this axis despite originating from different animals, plates, and analytical batches. Formal batch correction was <u>not</u> applied to this combined dataset. The ComBat^21^ algorithm assumes samples are drawn from a common distribution, and so it removes the between-group separation that is the object of this analysis.

**Figure 5:**
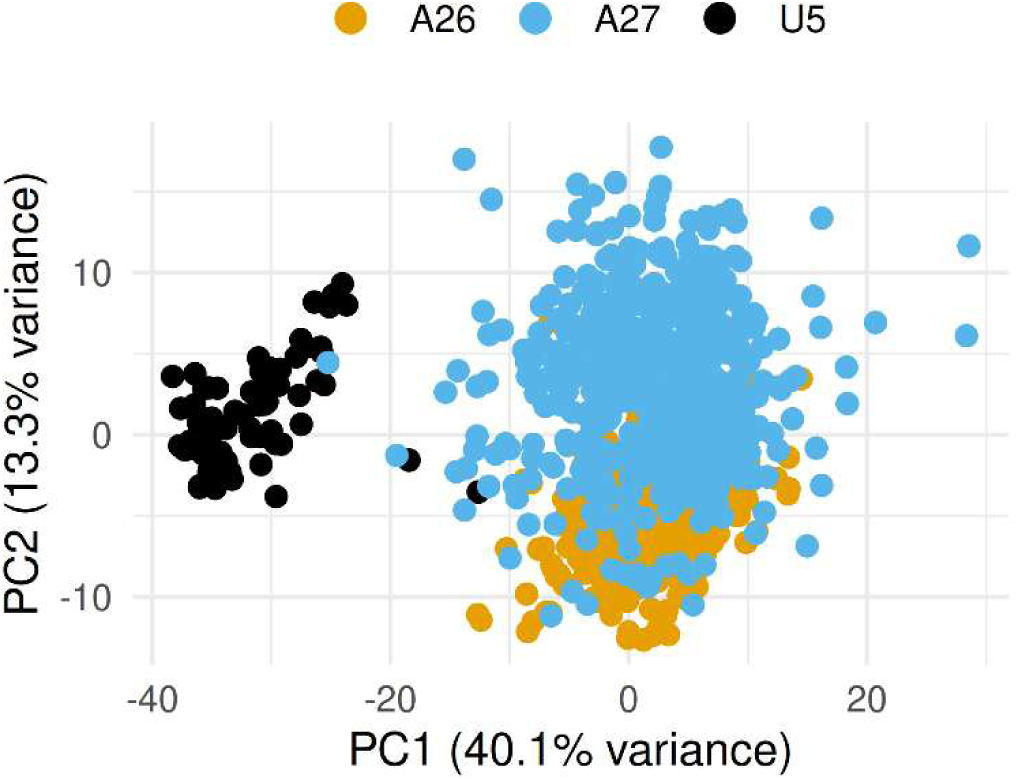
Principal Component Analysis (PCA) of metabolomic data with (U5) and without (A26, A27) blood. The dominant source of metabolomic variation when perfused and unperfused tissues are analyzed jointly is associated with perfusion/exsanguination status (separation of U5, black), exceeding variation attributable to inter-individual differences (separation of A26 and A27) or technical batch effects (within animal point dispersion for A26 and A27).

The metabolite classes loading positively on PC1 (enriched in exsanguinated tissue) were dominated by brain-parenchymal membrane lipids: phosphatidylcholines (both diacyl and ether-linked species), hexosylceramides, ceramides, and sphingomyelins, along with the amino acid derivatives SDMA and homoarginine. The metabolites loading negatively on PC1 (enriched in tissue that still contained blood) included triglycerides, diacylglycerols, spermine, and p-cresol sulfate—a gut-derived uremic toxin that circulates in blood plasma (see **Supplementary Table S2** for top 10 positive and negative loadings). This pattern is consistent with the expected effect of residual blood: unperfused tissue retains blood-borne lipids (triglycerides, diacylglycerols) and circulating metabolites (p-cresol sulfate), which dilute the brain-parenchymal lipid signal that dominates in exsanguinated tissue. The positive loadings were larger in magnitude (0.089–0.094) than the negative loadings (0.012–0.034), indicating that the PC1 separation is driven primarily by the unmasking of brain membrane lipid signatures when blood is removed, rather than by an influx of blood-derived analytes.

### Proteomic clusters based on PChclust are enriched with proteins associated with signaling and regulation

Clustering of the proteomic data by our novel PCA-based dimensionality reduction approach, PChclust (see Methods) identified 4717 clusters ranging from 1 to 696 proteins in each cluster. Eighty clusters containing at least 10 proteins were used for pathway enrichment analysis; we focus here on three that illustrate major identified biological pathways. **Supplementary Table S3** gives all 80 clusters.

#### Synaptic signaling, vesicle-mediated transport, neurotransmitters, cell-cell adhesion

This cluster contains 156 proteins with loadings ranging from -0.092 to -0.052. FDR significant pathways include regulation of synapse organization, modulation of chemical synaptic transmission, cell junction organization, and neuron projection development. The network analysis for this set of pathways shows a highly-structured graph with several hubs related to the neuronal processes (**Fig. 6a**; log fold changes: **Supplementary Fig. S6a**). Synaptic and post-synaptic pathways form a tight subnetwork at the lower right corner of the graph.

**Figure 6:**
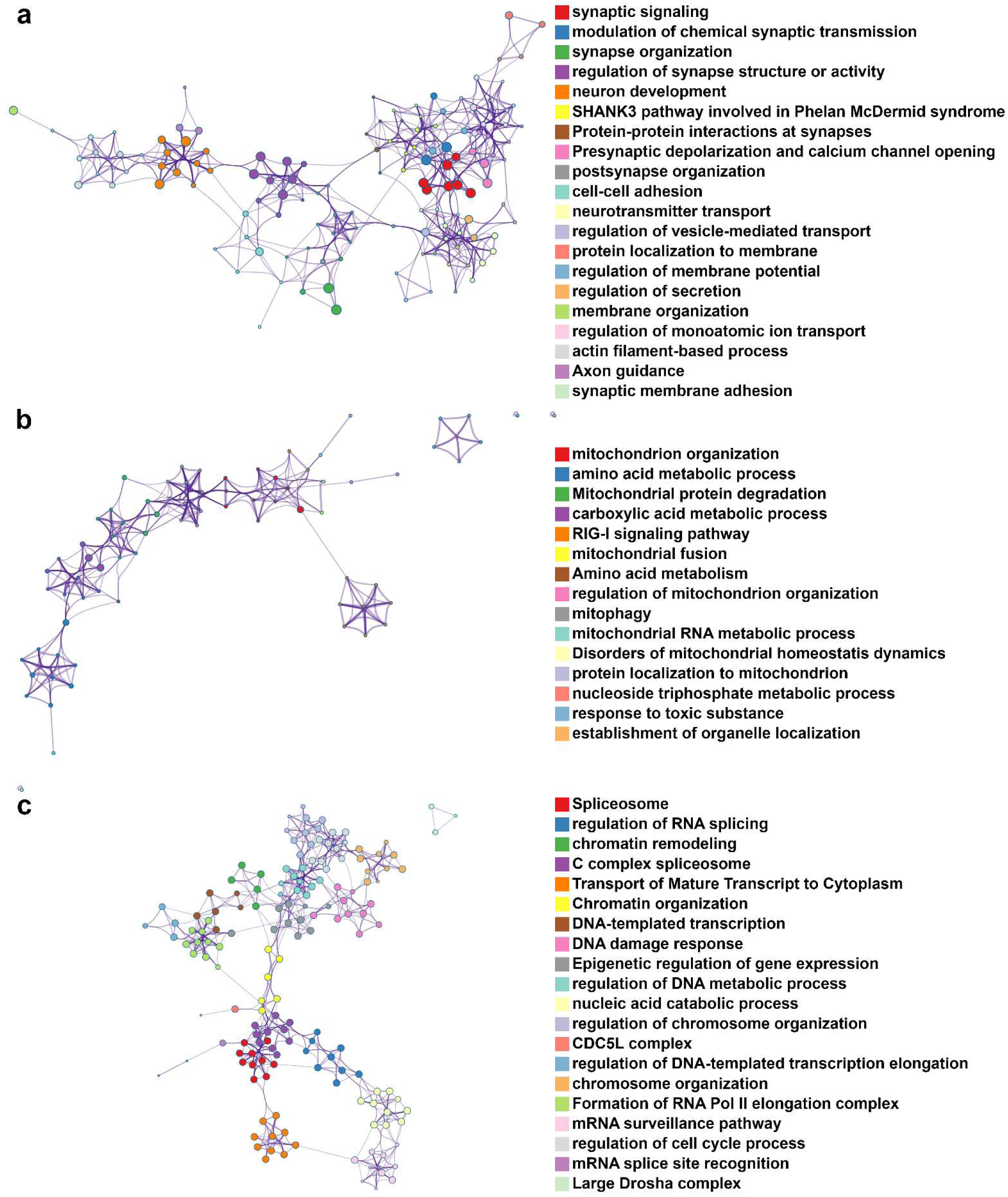
Biological pathways. Metascape^22^ network graphs for three protein clusters. **(a)** Synaptic signaling, vesicle-mediated transport, neurotransmitters, cell-cell adhesion. **(b)** Mitochondria, aerobic respiration. **(c)** Splicing.

#### Mitochondria, aerobic respiration

This cluster contains 54 proteins with loadings ranging from -0.152 to -0.120. FDR significant pathways include mitochondrial protein degradation, mitochondrion organization, carboxylic acid metabolic process, and mitochondrial fusion. The network of these pathways is loosely connected with one hub (**Fig. 6b**; log fold changes: **Supplementary Fig. S6b**).

#### Splicing

Splicing proteins are highly represented in one cluster of 696 proteins with loadings ranging from 0.024 to 0.044. FDR significant pathways include spliceosome, chromatin remodeling, regulation of RNA splicing, epigenetic regulation of gene expression, and other pathways related to splicing. The network graph is highly structured with several hubs evident (**Fig. 6c**; log fold changes: **Supplementary Fig. S6c**).

### Integrated pathway analysis reconstructs neuronal metabolic processes

To construct networks of both proteins and metabolites, we performed proteomics-driven joint pathway analysis (see Methods) for the three clusters described above (**Fig. 6**) against the KEGG annotation for Macaca mulatta, resulting in identification of comprehensive metabolic pathways.

#### Synaptic signaling, vesicle-mediated transport, neurotransmitters, cell-cell adhesion

Joint pathway analysis identified 220 pathways (33 FDR significant, p < 0.05; pathway impact: 0-0.56), including alanine, aspartate and glutamate metabolism, cysteine and methionine metabolism, and ABC transporters. The number of metabolites in the FDR significant pathways ranged from 2 to 27 while the number of proteins ranged from 0 to 6. ‘Neuroactive ligand-receptor interaction’ showed the optimal balance between proteins and metabolites identified in a pathway, with 9 metabolites and 6 proteins. Several synaptic pathways were identified, including synaptic vesicle cycle, GABAergic synapse, and glutamatergic synapse. **Table 2** shows the matched protein and metabolite features for the synaptic vesicle cycle pathway. Five enriched metabolites and six proteins were identified, covering both inhibitory (GABA, glycine) and excitatory (glutamic acid) signaling. The proteins are involved in synaptic signaling: calcium voltage-gated channel subunit alpha1 B (CACNA1B) is a presynaptic calcium channel, solute carrier family 6 member 7 (SLC6A7) is a transporter that removes L-proline from the extracellular space, vesicle-associated membrane protein 2 (VAMP2) and vesicular glutamate transporter 1 (SLC17A7) are synaptic vesicle proteins, and ATP6V0A1 and ATP6V0D1 are subunits of the vesicular proton pump.

**Table 2.**
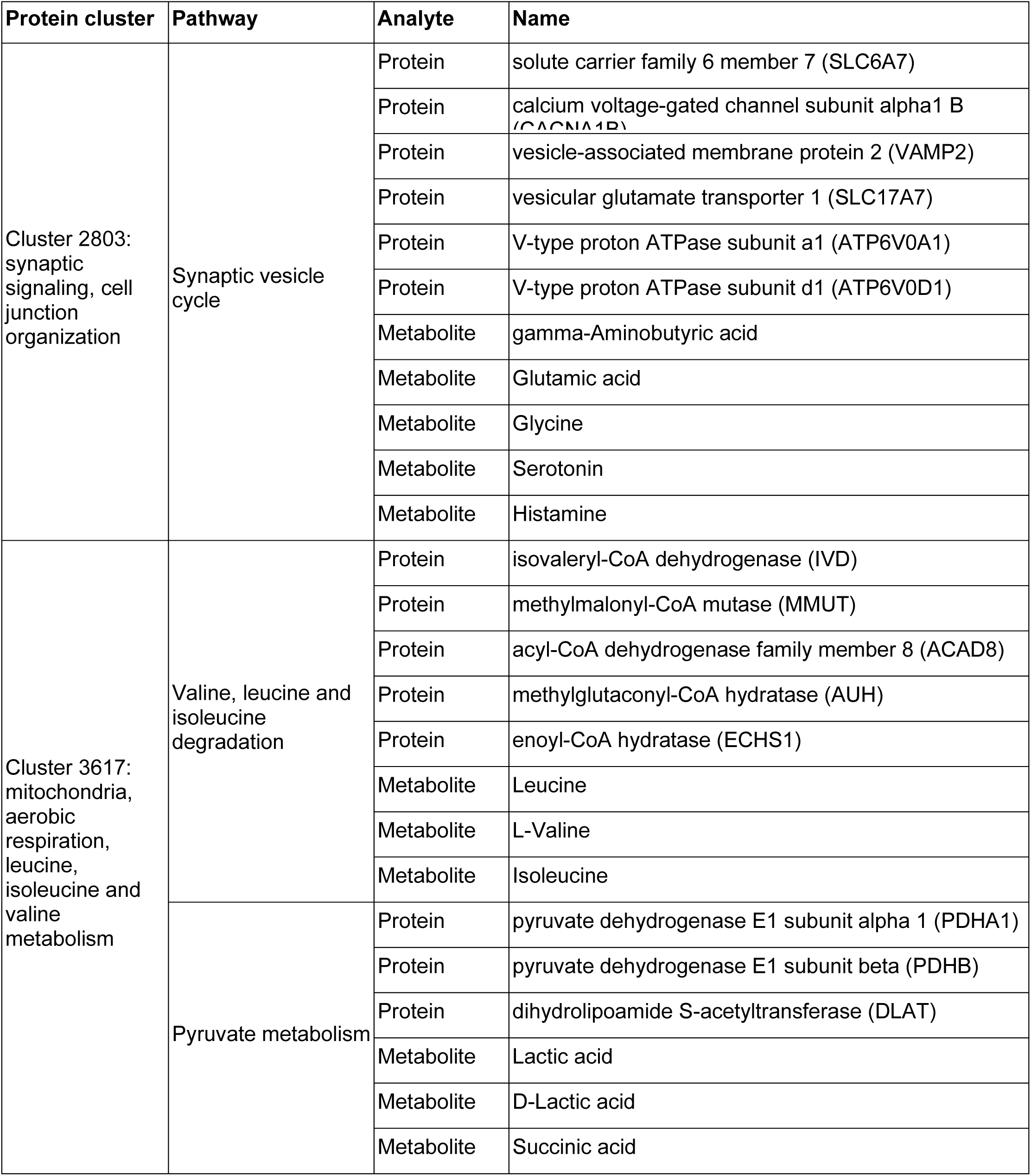
Matched protein and metabolite features from proteomics-driven joint pathway analysis.

| Protein cluster | Pathway | Analyte | Name |
|---|---|---|---|
| Cluster 2803: synaptic signaling, cell junction organization | Synaptic vesicle cycle | Protein | solute carrier family 6 member 7 (SLC6A7) |
|  |  | Protein | calcium voltage-gated channel subunit alpha1 B (CACNA1B) |
|  |  | Protein | vesicle-associated membrane protein 2 (VAMP2) |
|  |  | Protein | vesicular glutamate transporter 1 (SLC17A7) |
|  |  | Protein | V-type proton ATPase subunit a1 (ATP6V0A1) |
|  |  | Protein | V-type proton ATPase subunit d1 (ATP6V0D1) |
|  |  | Metabolite | gamma-Aminobutyric acid |
|  |  | Metabolite | Glutamic acid |
|  |  | Metabolite | Glycine |
|  |  | Metabolite | Serotonin |
|  |  | Metabolite | Histamine |
| Cluster 3617: mitochondria, aerobic respiration, leucine, isoleucine and valine metabolism | Valine, leucine and isoleucine degradation | Protein | isovaleryl-CoA dehydrogenase (IVD) |
|  |  | Protein | methylmalonyl-CoA mutase (MMUT) |
|  |  | Protein | acyl-CoA dehydrogenase family member 8 (ACAD8) |
|  |  | Protein | methylglutaconyl-CoA hydratase (AUH) |
|  |  | Protein | enoyl-CoA hydratase (ECHS1) |
|  |  | Metabolite | Leucine |
|  |  | Metabolite | L-Valine |
|  |  | Metabolite | Isoleucine |
|  | Pyruvate metabolism | Protein | pyruvate dehydrogenase E1 subunit alpha 1 (PDHA1) |
|  |  | Protein | pyruvate dehydrogenase E1 subunit beta (PDHB) |
|  |  | Protein | dihydrolipoamide S-acetyltransferase (DLAT) |
|  |  | Metabolite | Lactic acid |
|  |  | Metabolite | D-Lactic acid |
|  |  | Metabolite | Succinic acid |

#### Mitochondria, aerobic respiration

Joint pathway analysis identified 188 pathways (33 FDR significant, p < 0.05; impact: 0-0.86). Pathways covered included arginine, proline, alanine, aspartate, glutamate, serine, threonine, glutathione, and sphingolipid metabolism. The number of metabolites in the FDR significant pathways ranged from 1 to 27, and the number of proteins from 0 to 5. The valine, leucine, and isoleucine degradation pathway showed a balance between proteins and metabolites with 3 metabolites, 5 proteins. **Table 2** shows the matched protein and metabolite features for 1) valine, leucine, and isoleucine degradation: the proteins include dehydrogenases, hydratases, and a mutase, the metabolites include valine, leucine, and isoleucine; and 2) pyruvate metabolism, in which the three subunits of the pyruvate dehydrogenase complex (PDHA1, PDHB, DLAT) were identified along with 3 metabolites: lactic acid, D-lactic acid, and succinic acid.

#### Splicing

Joint pathway analysis identified 263 pathways (19 FDR significant, p < 0.05; impact: 0-1.0). Of the covered pathways, foremost was the spliceosome (p = 1.55·10^-59^). However, identified pathways were either entirely proteins—spliceosome (96 proteins), nucleocytoplasmic transport, ATP-dependent chromatin remodeling, and mRNA surveillance—or entirely metabolites: D-amino acid, aminoacyl-tRNA biosynthesis, and ABC transporters.

#### Integrated pathway analysis of the full proteomic and metabolomic results

For this analysis, all proteins and pathways were included. Loadings for the first PC for both proteins and metabolites were used as the scalar values (see Methods). The analysis identified 346 pathways (117 FDR, p < 0.05); pathway impact: 0-6.67). Major metabolic and signaling pathways relevant for neuronal function were identified as FDR significant, including endocytosis, axon guidance, glutamatergic synapse, sphingolipid signaling, and dopaminergic synapse. The number of metabolites in the FDR significant pathways ranged from 0 to 21 while the number of proteins ranged from 13 to 344. Among pathways with the highest number of metabolites, the efferocytosis pathway matched to 7 metabolites and 140 proteins; synaptic vesicle cycle mapped to 5 metabolites and 74 proteins.

Interestingly, neurodegenerative diseases had a large number of mapped proteins including Alzheimer’s disease (271 proteins) and Parkinson’s disease (200). The MAPK signaling pathway mapped to 1 metabolite and 191 proteins. Pathways that showed the largest number of metabolites and proteins (noting that the number of proteins in the dataset >> the number of metabolites) included synaptic vesicle cycle (5 metabolites, 63 proteins) and sphingolipid signaling (4 metabolites, 94 proteins).

## Discussion

Here, we present a framework for generating spatially registered, paired metabolomic and proteomic maps across an entire cortical hemisphere in the rhesus monkey, together with analysis tools that explicitly incorporate spatial dependence. The core design decision behind our work was to have assays paired at the sample level—each voxel having been split to yield matched aliquots—rather than integrated by post hoc alignment across different dissections or animals. To handle dimensionality and collinearity, we developed PChclust, a principal-component–guided feature clustering method that can be applied to samples within each animal separately to mitigate collinearity among high-dimensional features without assumptions on the dependency across samples. For cross-omic integration, we developed sr-sCCA, which incorporates spatial neighborhood structure rather than assuming sample independence. The loadings obtained from sr-sCCA and PChclust can be used to obtain weights for original features (i.e., analytes) for each animal, which can then be compared and transferred between animals. Similar to traditional hierarchical clustering and canonical correlation analysis, PChclust and sr-sCCA do not carry the strict sample size requirements typical of hypothesis-testing frameworks. While low sample sizes traditionally risk overfitting in high-dimensional spaces, the integration of collinearity reduction by PChclust, alongside spatial and sparsity regularizations in sr-sCCA, stabilizes the latent structure extraction even with limited samples. Importantly, as we apply these methods, the sample size is tissue voxels (hundreds of samples), not animals (two). Our approach to integrated pathway analysis — using proteomic clusters to organize the joint analysis — follows a rationale shared by prior multi-omics studies^23, 26^: the proteome, with its far greater dimensionality, provides the structural scaffold, while metabolites supply functional readout (for comprehensive reviews of multi-omics integration strategies and approaches, see^17,26–32^).

Several lines of evidence indicate that our analysis of the resulting data captures real biology at the spatial scale of our measurements, and measurement scale itself was a key variable in our work. Both sampling resolutions (∼4 mm for A26, ∼2.5 mm for A27) captured spatial decay of molecular similarity, but the denser sampling in A27 resolved regional heterogeneity (particularly in the metabolome) that was not detectable in A26.

There is a cost-resolution tradeoff in this approach that scales with voxel number. Our method is advantageous when studying patterns at the millimeter-to-centimeter scale across a structure too large for imaging-based platforms. Where only a few regions are of interest, or sub-millimeter resolution is essential, conventional dissection-based profiling or imaging methods are likely to be more cost-effective. The neighbor-correlation analysis described above (Fig. 4) offers a practical entry point to our approach: an initial survey identifies where molecular profiles are spatially correlated, and so can be sampled sparsely, and where they are not, allowing sampling effort, and cost, to be concentrated where finer resolution is informative. Importantly, at both the resolutions we tested here, sr-sCCA identified joint proteome–metabolome components that were spatially coherent, and cross-animal projection, i.e. using the feature weights from A27 to project feature measurements from A26 revealed conserved spatial structure (i.e., a consistency in the underlying spatial gradients of measured analytes).

### Non-uniqueness of the decomposition

As with other high-dimensional latent decompositions, the sr-sCCA solution is not, in general, uniquely identified. When two canonical correlations are close, individual vectors within that near-degenerate subspace are not anchored, although the subspace itself is well determined. Under strong collinearity, the choice of which among several correlated analytes the sparsity penalty retains is unstable. In our approach, these indeterminacies concentrate in the feature loadings, with the spatial score maps being the more stable output. Traditional CCA is the opposite: high-dimensional collinearity destabilizes loadings and overfits scores, often yielding mathematically perfect but biologically meaningless correlations. Because PChclust scores represent the mapped data and remain geometrically stable, applying it first mitigates the collinearity, so the downstream sr-sCCA is more stable. It is still not guaranteed to yield a single, unique mathematical solution; instead, our spatial regularization and sparsity constraints work together to guide the optimization away from noisy, arbitrary solutions and toward a stable, biologically meaningful one. Our analysis framework is partly self-constraining in that the spatial penalty selects among otherwise-equivalent solutions in favor of spatially coherent ones, and PChclust collapses collinear features so that pathway-level readouts are robust to the interchange of individual analytes within a correlated set.

This delimits where the method is appropriate without modification. It is well suited to detecting and comparing spatial gradients, boundaries, and regional patterns, and to assessing whether such patterns are conserved across samples or animals, because these are read from the scores. It is not, without further constraint, suited to claims about the identity of individual molecular markers. Similarly, with regard to underlying biochemistry, interpretation is most secure at the cluster or pathway, rather than individual analyte, level. Integration across the proteomic and metabolomic results further increases confidence for specific clusters and pathways. Cross-sample or cross-animal projection, as used here, provides a practical filter for retaining components with reproducible spatial structure. Where molecular identity is of interest, stability selection—refitting across resamples and initializations and reporting selection frequencies—is advisable; because neighboring voxels are correlated, this resampling should draw spatially contiguous blocks rather than independent voxels.

### Identifying biochemical pathways

A separate pathway enrichment analysis on the output of PChclust recovered major neuronal processes — synaptic signaling, mitochondrial respiration, RNA splicing — and integrated pathway analysis reconstructed complete metabolic circuits within single voxels. Our recovery of neurotransmitter pathways (GABA, glycine, glutamate, alongside CACNA1B and SLC6A7) from an unsupervised spatial decomposition of data from bulk homogenized cortical tissue provides biological face validity. Finally, that neurodegenerative disease pathways — including Alzheimer’s disease (AD; 271 proteins) and Parkinson’s disease (200 proteins) — emerged in our analysis underscores that the molecular infrastructure implicated in these diseases is detectable by our framework. Further study is needed to understand whether disease-associated vulnerability is spatially organized, or transfers across individuals.

### Comparison with other spatial multi-omics approaches

Current spatial molecular methods (such as mass spectrometry imaging, spatial transcriptomics, multiplexed protein imaging) offer micrometer-scale, in some cases subcellular, resolution, but they face a fundamental spatial coverage problem when applied to large brains. Even with spatial tiling, current imaging platforms are insufficient to map a primate cortical hemisphere at high resolution. Our approach sacrifices resolution (millimeters rather than micrometers) and gains whole-hemisphere coverage and deep molecular profiling (∼10,000 protein groups and ∼330 metabolites per voxel). We cannot resolve laminar, columnar, or single-cell structure, but we can map gradients, boundaries, and regional patterns across the full spatial extent of cortex. For the class of questions that motivate our work (vulnerability and resilience in aging and Alzheimer’s disease; AD), relevant spatial patterns of pathology span centimeters, and so this trade-off seems reasonable.

### Why non-human primates, and why exsanguination matters

Spatial patterns of pathology in AD have fine-grained features across association cortex; territory for which rodent models offer limited homology. Furthermore, rodents have no endogenous genotype that yields an AD-like aging trajectory, and transgenic mouse models, designed to aggressively generate pathology, almost certainly do not recapitulate the complex interplay of triggers and tissue responses that characterizes early aging and preclinical AD. Dense spatial mapping of the kind presented here would be difficult to obtain from humans directly; analyzing a whole human cortical hemisphere represents a significant opportunity cost that may not be justifiable. Furthermore, preservation and isolation of biomolecules in human samples is challenging.

In our comparison with unperfused tissue, exsanguination-associated variance dominated the metabolomic data structure (PC1 ∼40%), driven by blood-borne lipids and circulating metabolites rather than brain-parenchymal species. The metabolites that separated the exsanguinated (A26 and A27) versus not exsanguinated (U5) tissue were not major contributors to any sr-sCCA component, so residual blood is unlikely to account for the spatial patterns we observed. This does not, however, guarantee that equivalent analyses on tissue that has not been exsanguinated would recover the same components — blood-associated metabolites could share spatial structure with brain-parenchymal species in a CCA decomposition, yielding components that reflect a different mixture of biological and artifactual variance. The macaque, which can offer controlled harvest conditions, enables a class of spatially resolved biochemical studies that are, at best, hard to do well in humans.

### Limitations

The most obvious limitation of this study is sample size. While this is primarily a methods-development report (not designed for inference), with the metabolome and proteome coming from the same homogenate, and maps being defined across each animal’s own cortex, the hundreds of voxels per animal are the biological replicates. The cross-animal projection analysis provides evidence of conserved spatial patterns. Population-level inference will nonetheless require additional cases, particularly to address aging trajectory and biological sex.

The targeted metabolomics panel constrains coverage to ∼330 metabolites after quality filtering; untargeted metabolomics would expand coverage, but also lose the quantitative standardization that facilitates cross-plate and cross-animal comparison. Our voxel sizes (∼2.5–4 mm) do not support laminar or columnar resolution, being designed for area-level comparison and gradient detection. Our tissue handling timeline for hand dissection (∼2.5 hours from anoxia to last frozen voxel) introduces the possibility of metabolite degradation in later-sampled voxels; we counterbalanced dissection order across animals and did not observe opposing spatial gradients.

The exsanguination comparison was performed using available tissue and has its own limitations: U5 differed from A26 and A27 in age (∼2.5 vs. 22–24 years), sex (male vs. female), tissue handling, and laboratory of origin. Had age or sex been the dominant factor in the PCA separation, we would have expected contributions from hormones or energy metabolites; instead, loadings were dominated by phosphatidylcholines, hexosylceramides, triglycerides, and circulating metabolites consistent with the presence or absence of blood. U5 tissue was flash-frozen within minutes, while A26 and A27 voxels were frozen over ∼2.5 hours; labile metabolites — particularly acylcarnitines — may have been better preserved in U5. However, for most of the metabolites driving the PC1 separation, the direction of the difference is inconsistent with a preservation artifact: the species enriched in U5 were predominantly blood-borne (triglycerides, p-cresol sulfate), and the species enriched in A26/A27 were intact membrane lipids that degradation would consume, rather than create.

### What our method enables

We do not yet know at what spatial scales molecular vulnerability and resilience patterns are organized in the aging primate cortex; whether they are gradients spanning lobes, patchy territories organized by cytoarchitecture, or fine-grained mosaics following local connectivity. The answer matters for both mechanism and intervention, because it will determine what measurements would be needed to detect early change, and what spatial resolution would be required for targeted therapies. The data presented here cannot answer these questions; we studied two healthy animals. However, at 22 and 24 years of age (human-equivalent ∼60-70 years), both animals are squarely within the window corresponding to the human LOAD prodrome and their unremarkable biomarker profiles are consistent with that staging; the prodrome, by definition, precedes detectable pathology. Furthermore, the framework we have developed makes it feasible to ask them across a series of animals spanning the aging trajectory, with paired molecular and histopathological atlases. The methods — voxelation with paired multi-omic profiling, spatially aware dimensionality reduction, and cross-omic integration that respects spatial structure — are our contributions. The insights will come from applying them.

## Acknowledgements and Author Contributions

The authors gratefully acknowledge T. Nguyen for surgical assistance and N. Schiff for assistance with reconstruction of voxelated cortical sheets. We also thank the Duke Proteomics and Metabolomics Core Facility for generating the metabolomic and proteomic data, including Matthew Foster, Lucas Li, Marlene Violette, Sophia Guerrero, Greg Waitt, and Lisa St. John-Williams.

AAD conceived of, designed and supervised the study. AAD and AMB performed tissue harvest surgeries. AMB prepared samples, including blinding, and reconstructed voxelated cortical sheets. PS devised the PChclust and sr-sCCA statistical methods, with QW implementing them. MWL performed the pathway enrichment and transomic integration analyses. AAD wrote the manuscript. All authors reviewed the manuscript.

## Data and Code Availability

Processed proteomics data: https://doi.org/10.7924/r4r506

Metabolomics data and sample-position tables: https://doi.org/10.5281/zenodo.23857927

Analysis code https://doi.org/10.5281/zenodo.23868034

