## Supplementary Information, Tables and Figures for "Integrated metabolomics and proteomics from voxelated cortical hemispheres of adult rhesus monkeys": omics_SuppFigs.pdf

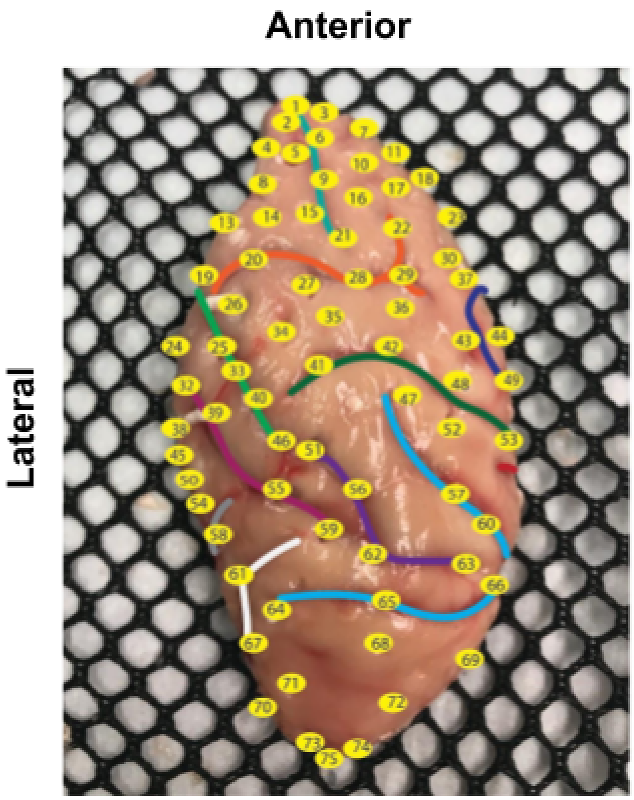


**Supplementary Figure S1: Tissue punch map, U5.** Dorsal surface, left hemisphere (thawed) after punch sampling. Numbered yellow ovals: punch locations. Colored lines: sulci.

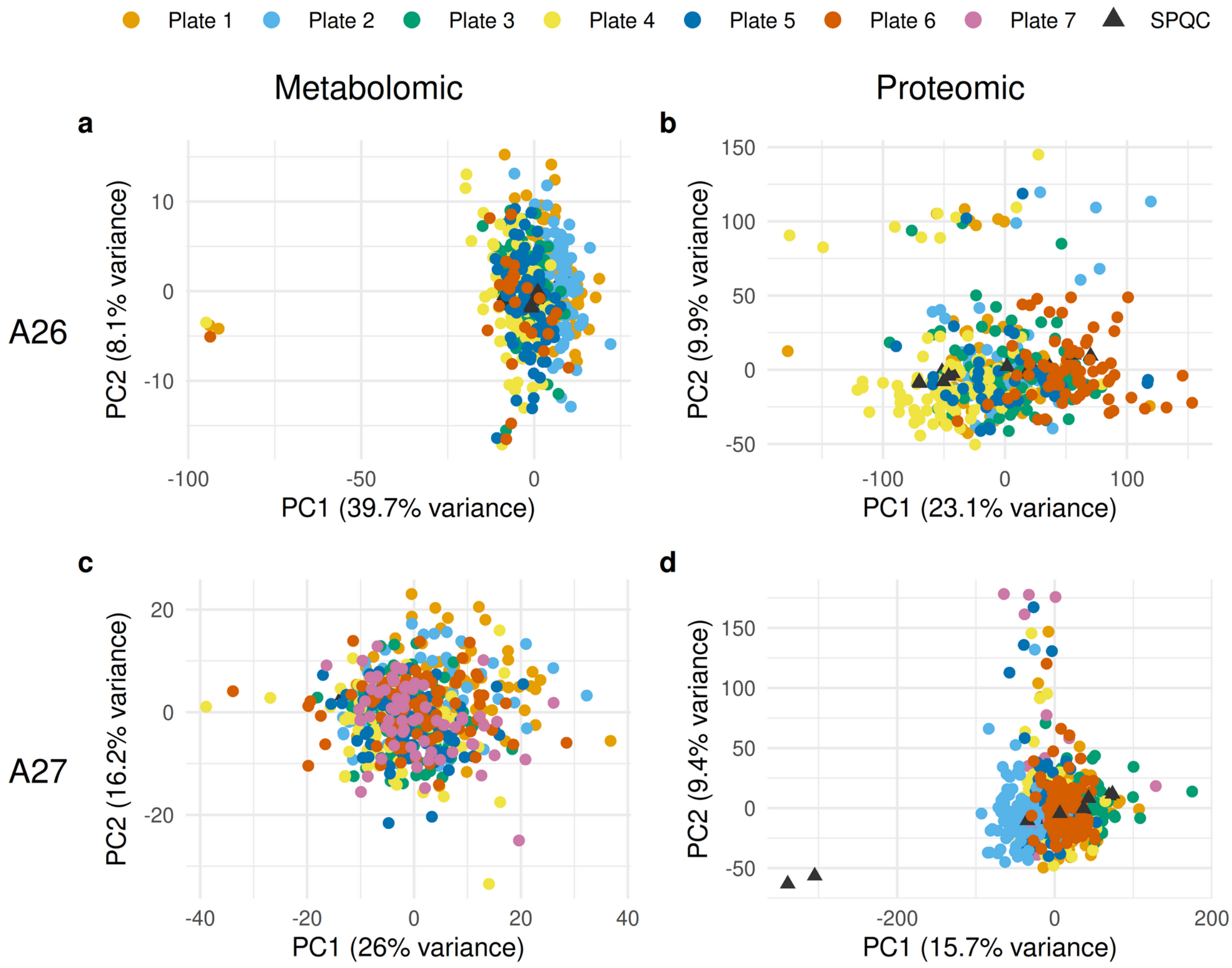


**Supplementary Figure S2: Pre-ComBat PCA plots.** (A) A26 metabolomics. (B) A26 proteomics. (C) A27 metabolomics. (D) A27 proteomics. Points are colored by processing plate; black triangles indicate SPQC replicates. Plate-driven separation visible here, particularly in the metabolomics data for A26, was substantially reduced by ComBat correction (compare to Figure 2).

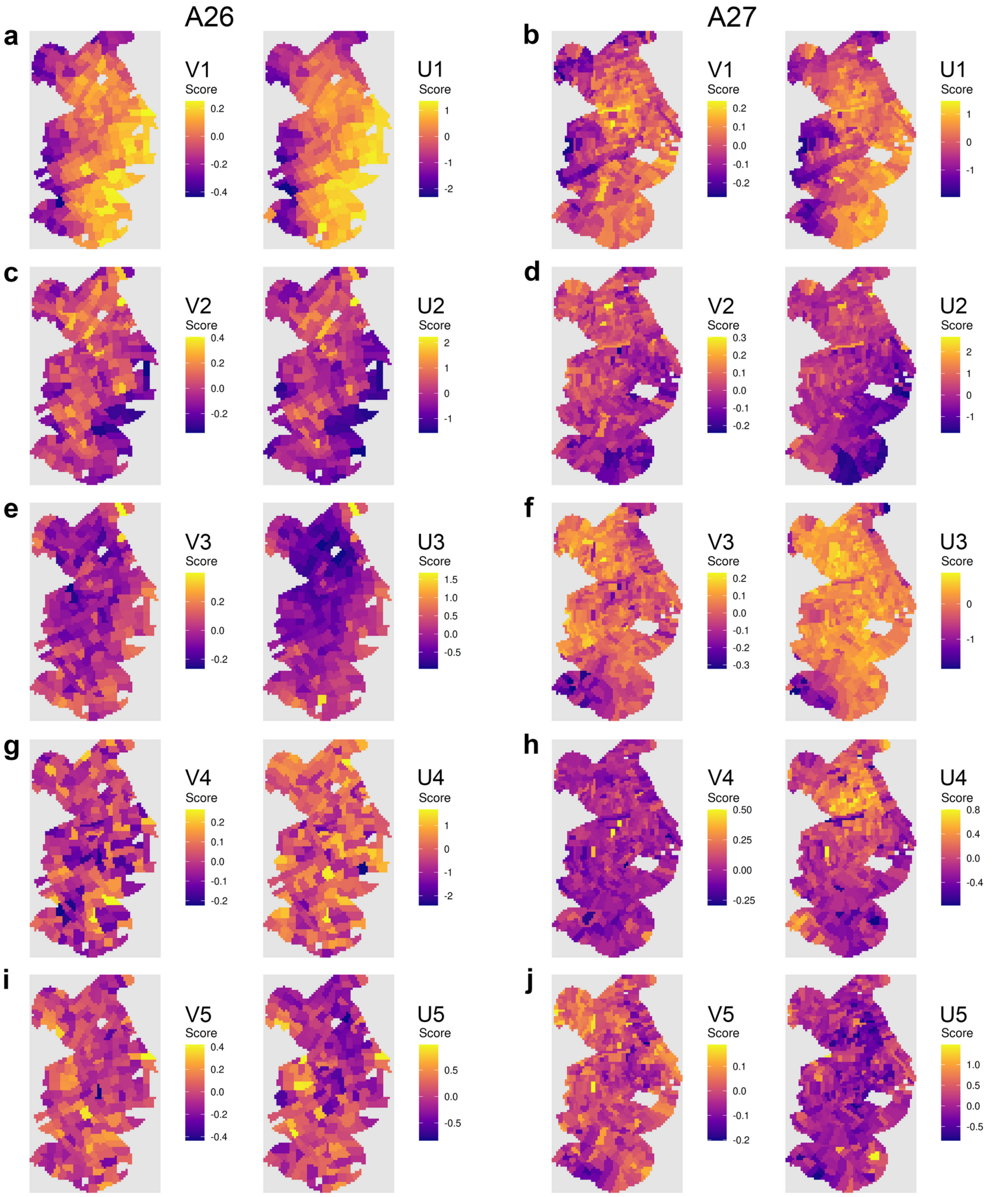


**Supplementary Figure S3: Spatial maps for sr-sCCA components 1-5.** Metabolomic canonical scores (V) and proteomic canonical scores (U) for components 1–5, projected onto cortical spatial coordinates for A26 (left) and A27 (right). Color scales are independently normalized within each map. Warm colors: high/positive scores; cool colors: low/negative scores. Components 1 and 2 are also shown in Figure 3 of the main text.

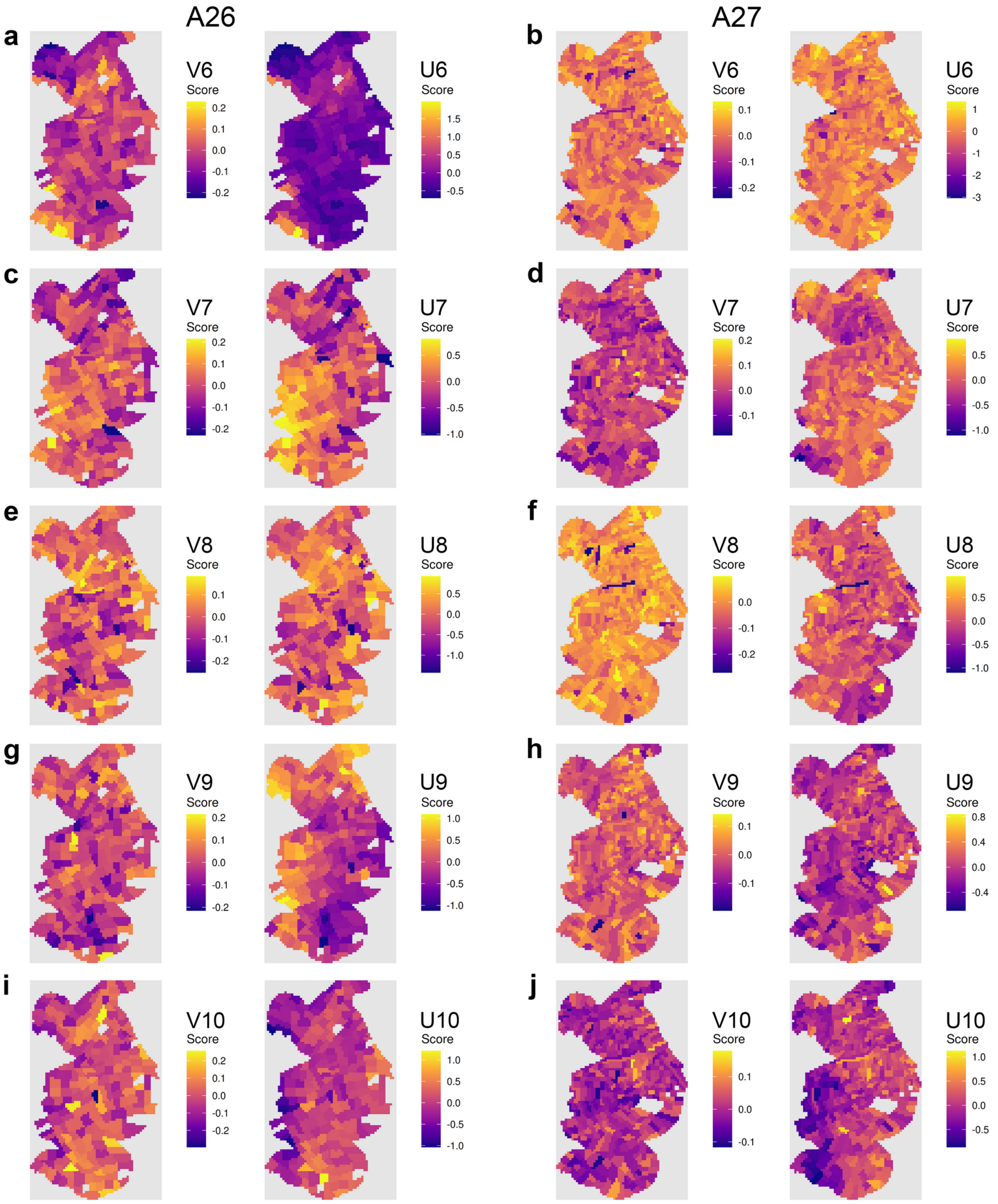


**Supplementary Figure S4: Spatial maps sr-sCCA components 6-10.** Metabolomic canonical scores (V) and proteomic canonical scores (U) for components 6–10, projected onto cortical spatial coordinates for A26 (left) and A27 (right). Color scales are independently normalized within each map. Warm colors: high/positive scores; cool colors: low/negative scores.

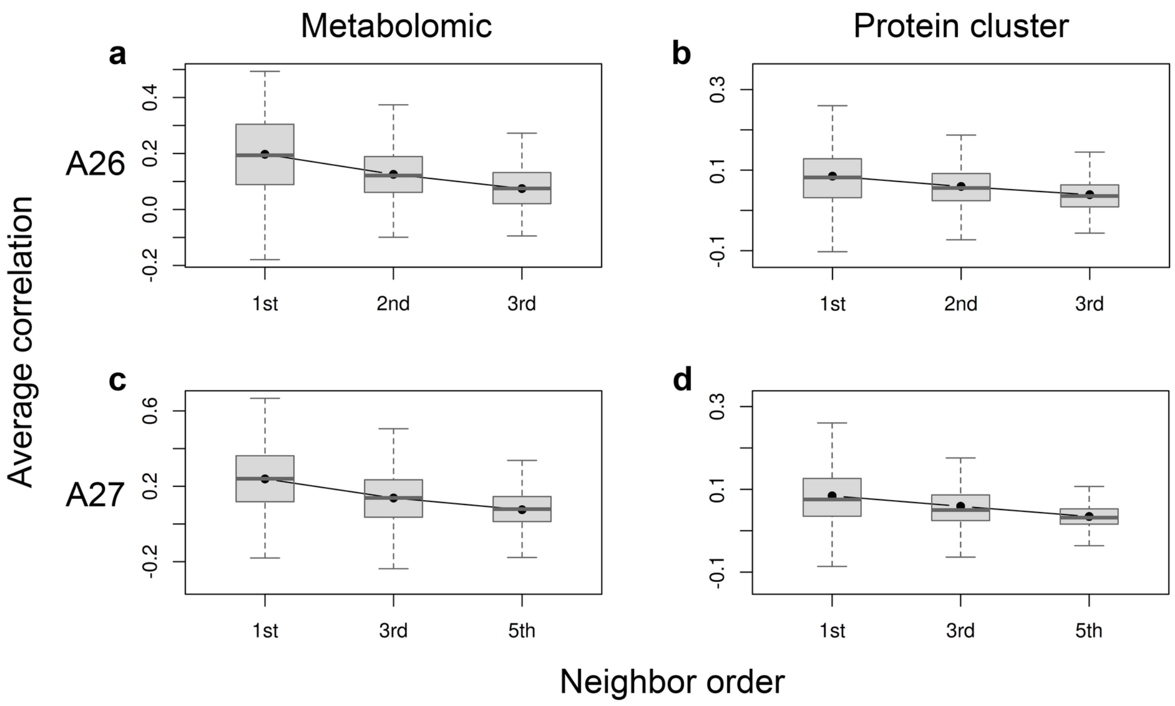


**Supplementary Figure S5: Spatial gradient boxplots.** Mean correlations and their standard errors were overlaid to illustrate the trend of molecular similarity across increasing spatial distance.

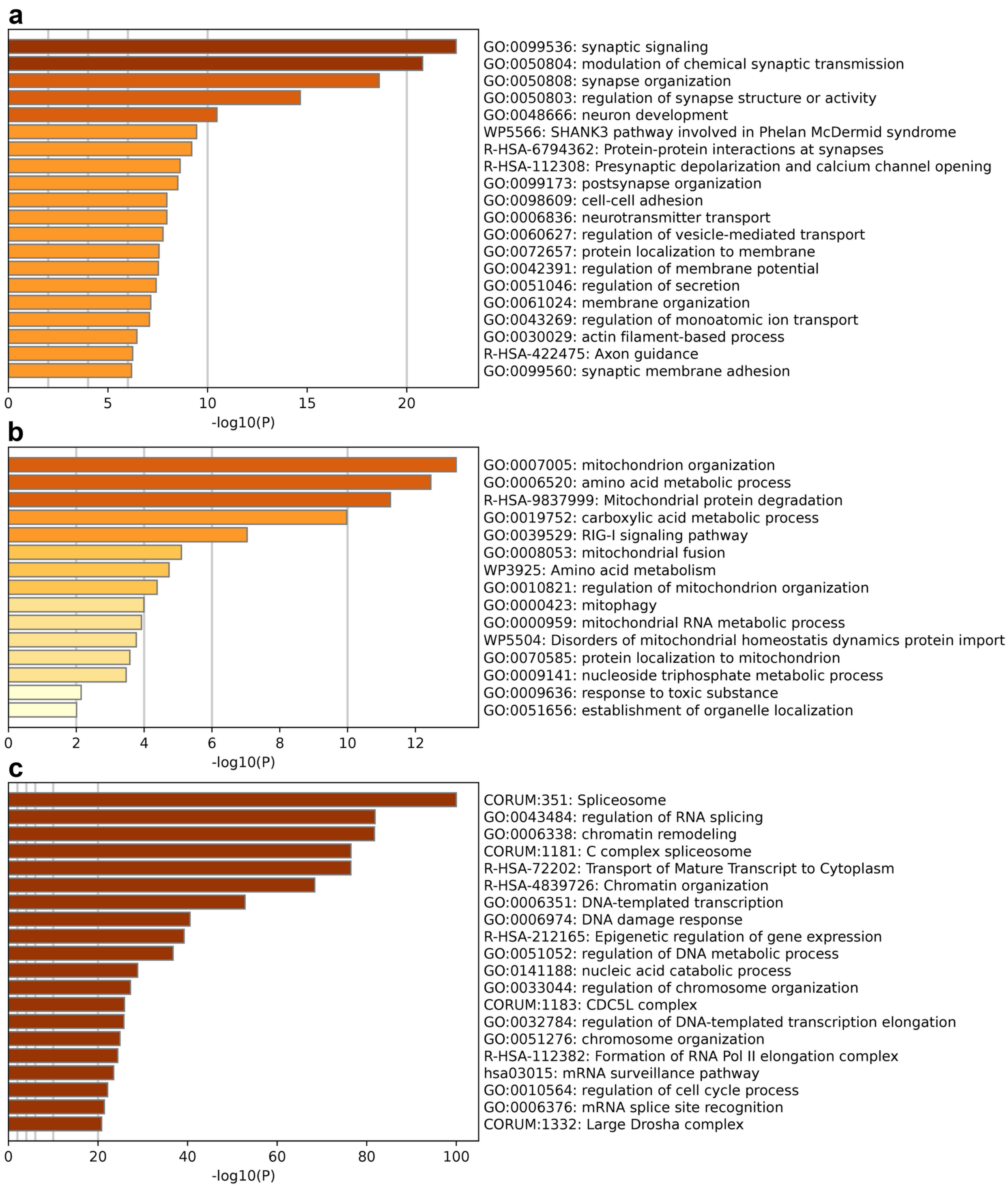


**Supplementary Figure S6: Log fold change plots from pathway analysis for three proteomic clusters.** (a) Synaptic signaling, vesicle-mediated transport, neurotransmitters, cell-cell adhesion. (b) Mitochondria, aerobic respiration. (c) Splicing.
