## Supplementary Information, Tables and Figures for "Integrated metabolomics and proteomics from voxelated cortical hemispheres of adult rhesus monkeys": omics_SuppInfo.pdf

### Supplementary Information: Animal Health Histories

All animal procedures prior to the studies described in the manuscript associated with this Supplement were conducted at the Oregon National Primate Research Center (ONPRC) at Oregon Health & Science University (OHSU). Both animals were adult female Indian-origin rhesus macaques (Macaca mulatta) that were born at the ONPRC and had been housed there since birth. Prior to the present study, both animals had been assigned to one or more IACUC-approved research protocols and had undergone the procedures summarized below. Full medical, dietary, social housing, and health records from birth are available upon request.

#### A26 (USDA #21545)

**Demographics.** A26 ‘Bonnie’ was a female rhesus macaque born on April 20, 2000 via vaginal delivery at the ONPRC outdoor breeding colony (Corral 2). At the time of tissue collection for the present study, she was approximately 24 years of age. Her last recorded body weight prior to tissue collection was 8.8 kg (May 2023) with a body condition score (BCS) of 3/5.

**Prior research participation.** A26's research history included participation in a timed-mated breeding (TMB) program and assignment to a maternal-fetal physiology protocol that involved intrauterine catheterization, serial amniocentesis, fetal MRI, and delivery by cesarean section. She underwent two cesarean sections in total. She was subsequently assigned to a protocol involving chronic low-dose ethanol administration (0.5 g/kg/day) with operant conditioning, during which she was singly housed. She also underwent a single bronchoalveolar lavage (BAL) procedure in August 2020 as a day-lease to a separate protocol. Brain MRI scans were conducted during her maternal-fetal protocol and again in January 2021 and April 2022 as part of the present study.

**Significant medical history.** A26 had a documented history of chronically elevated serum alanine aminotransferase (ALT) in the range of 100–250 IU/L, with associated mild hepatomegaly noted on physical examination. This was managed with chronic ursodiol supplementation, to which the ALT consistently responded, and was monitored at semiannual physical examinations. A comprehensive chemistry panel in October 2022 showed ALT within the normal reference range while on ursodiol. She was also diagnosed with moderate degenerative joint disease (DJD) affecting primarily the hips (moderate decreased range of motion, left greater than right), with milder involvement of the shoulders, elbows, and stifles. This was managed with daily oral glucosamine/chondroitin supplementation. She had a history of chronic pelleted stool managed with daily oral psyllium and periodic prunes, and dorsal xerosis (a dry, flaky skin patch at the thoracolumbar junction) managed with periodic medicated chlorhexidine baths. Mild to moderate dental tartar, gingivitis, and age-related dental wear were noted at physical examinations. She also had a single resolved corneal ulcer (right eye, January 2021) and a resolved episode of colitis (April 2016).

**Menstrual history and reproductive status.** A26 menstruated regularly throughout the period covered by the clinical record, with menses documented at approximately monthly intervals through April 2023, consistent with continued ovarian cyclicity. Serum FSH and LH levels were assayed from a blood sample collected on April 5, 2024, when the animal was 23.9 years of age (see **Supplementary Table S1**). FSH was 0.98 ng/mL (mean of duplicates; CV 12.7%) and LH was 4.74 pg/mL (mean of duplicates; CV 19.8%). These values are within the pre-menopausal range for female rhesus macaques and, together with the regular menstrual record, indicate that A26 had not undergone menopause at the time of sampling.

#### A27 (USDA #22926)

**Demographics.** A27 ‘Ms. Frizzle’ was a female rhesus macaque born on April 1, 2002 via vaginal delivery at the ONPRC outdoor breeding colony (Animal Run 3). At the time of tissue collection for the present study, she was approximately 22 years of age. Her last recorded body weight prior to tissue collection was 6.8 kg (April 2023) with a BCS of 3/5.

**Prior research participation.** A27's research history included participation in a timed-mated breeding program and assignment to a maternal-fetal physiology protocol that involved fetal and maternal vascular catheterization (femoral artery and vein), intrauterine catheterization, serial amniocentesis, and delivery by cesarean section. She underwent one cesarean section in total. She also had a gastrocnemius muscle biopsy performed at the time of cesarean delivery. Her research assignment history included a glucose tolerance test and serial amniotic fluid sampling during the catheterization protocol.

**Significant medical history.** A27 had a documented history of mild to moderate degenerative joint disease affecting the hips (right greater than left), treated with daily oral glucosamine/chondroitin supplementation beginning in late 2021. Mild to moderate dental tartar, calculus, and gingivitis were noted, and she received a dental prophylaxis in November 2022. Three small (1–1.5 cm) subcutaneous masses on the ventral midline abdomen (presumed benign lipomas) and a small (3–5 mm) smooth, firm nodule at the dorsal left cervix/distal uterus were noted at physical examinations and monitored. She had two brief episodes of diarrhea (February 2016; July 2021); both resolved rapidly with bismuth subsalicylate treatment. A27 sustained a number of injuries secondary to low social status and outdoor social housing, including a crush trauma presenting with likely rhabdomyolysis (October 2014), hand lacerations (December 2014), heel laceration (July 2015), and a tail degloving (August 2019). A27 was on active weight management for much of her adult life, with target weight ranges adjusted over time.

**Menstrual history and reproductive status.** A27 menstruated regularly throughout the period covered by the clinical record, with menses documented at approximately monthly intervals through April 2023, consistent with continued ovarian cyclicity. Serum FSH and LH levels were assayed from a blood sample collected on May 29, 2024, when the animal was 22.2 years of age (see **Supplementary Table S1**). FSH was 0.96 ng/mL (mean of duplicates; CV 4.2%) and LH was 5.64 pg/mL (mean of duplicates; CV 9.7%). As with A26, these values fall within the pre-menopausal range for female rhesus macaques and, together with the regular menstrual record, indicate that A27 had not undergone menopause at the time of sampling.
