## Supplementary Information, Tables and Figures for "Integrated metabolomics and proteomics from voxelated cortical hemispheres of adult rhesus monkeys": omics_SuppTables1and2.pdf

**Supplementary Table S1. Serum gonadotropin levels**

*Single serum samples were assayed for follicle-stimulating hormone (FSH) and luteinizing hormone (LH) to assess reproductive/menopausal status. Assays were performed by the Biomarkers Core Laboratory at the Emory National Primate Research Center (supported by NIH P51 OD011132).*

| Animal | Age at Sample | Sample Date | FSH (ng/mL) | LH (pg/mL) | Interpretation |
|---|---|---|---|---|---|
| A26 (#21545) | 23.9 yr | 4/5/2024 | 0.98 | 4.74 | Pre-menopausal |
| A27 (#22926) | 22.2 yr | 5/29/2024 | 0.96 | 5.64 | Pre-menopausal |

***Note:*** *In female rhesus macaques, menopause typically occurs between approximately 25 and 27 years of age and is characterized by sustained elevation of serum FSH (typically > 5–10 ng/mL, depending on the assay) accompanied by cessation of menses. The low FSH concentrations observed here (~1 ng/mL for both animals), combined with documented regular menstrual cyclicity through 2023, are consistent with pre-menopausal reproductive status at the time of sampling.*

**Supplementary Table S2. Top ten positive and negative features driving PC1**

| Feature | PC1 Loading | PC1 Loading Absolute Value |
|---|---|---|
| PC aa C36:1 | 0.094 | 0.094 |
| PC aa C42:5 | 0.093 | 0.093 |
| PC ae C38:5 | 0.092 | 0.092 |
| PC ae C38:4 | 0.092 | 0.092 |
| PC aa C34:1 | 0.091 | 0.091 |
| PC ae C34:1 | 0.091 | 0.091 |
| PC ae C36:4 | 0.091 | 0.091 |
| PC ae C38:6 | 0.091 | 0.091 |
| HexCer (d18:1/24:1) | 0.091 | 0.091 |
| PC ae C42:2 | 0.090 | 0.090 |
| PC aa C36:0 | -0.034 | 0.034 |
| TG (22:6_32:0) | -0.031 | 0.031 |
| Spermine | -0.026 | 0.026 |
| TG (20:4_34:1) | -0.024 | 0.024 |
| DG (14:1_18:1) | -0.023 | 0.023 |
| TG (20:4_36:4) | -0.022 | 0.022 |
| TG (18:1_38:5) | -0.019 | 0.019 |
| TG (18:1_38:6) | -0.019 | 0.019 |
| TG (22:6_34:1) | -0.017 | 0.017 |
| Cer (d18:2/16:0) | -0.012 | 0.012 |
